# Folding RNA in mixtures of monovalent and divalent cations: Theory and simulations

**DOI:** 10.1101/732917

**Authors:** Hung T. Nguyen, Naoto Hori, D. Thirumalai

## Abstract

RNA molecules cannot fold in the absence of counter ions. Experiments are typically performed in the presence of monovalent and divalent cations. How to treat the impact of a solution containing a mixture of both ion types on RNA folding has remained a challenging problem for decades. By exploiting the large concentration difference between divalent and monovalent ions used in experiments, we develop a theory based on the Reference Interaction Site Model (RISM), which allows us to treat divalent cations explicitly while keeping the implicit screening effect due to monovalent ions. Our theory captures both the inner shell and outer shell coordination of divalent cations to phosphate groups, which we demonstrate is crucial in an accurate calculation of RNA folding thermodynamics. The RISM theory for ion-phosphate interactions when combined with simulations based on a transferable coarse-grained model allows us to accurately predict the folding of several RNA molecules in a mixture containing monovalent and divalent ions. The calculated folding free energies and ion-preferential coefficients for RNA molecules (pseudoknots, a fragment of the ribosomal RNA, and the aptamer domain of the adenine riboswitch) are in excellent agreement with experiments over a wide range of monovalent and divalent ion concentrations. Because the theory is general, it can be readily used to investigate ion and sequence effects on DNA properties.

**Significance Statement:** RNA molecules require ions to fold. The problem of how ions of differing sizes and valences drive the folding of RNA molecules is unsolved. Here, we take a major step in its solution by creating a method, based on the theory of polyatomic liquids, to calculate the potential between divalent ions and the phosphate groups. The resulting model, accounting for inner and outer sphere coordination of Mg^2+^ and Ca^2+^ to phosphates, when used in coarse-grained molecular simulations predicts folding free energies for a number of RNA molecules in the presence of both divalent and monovalent ions that are in excellent agreement with experiments. The work sets the stage for probing sequence and ion effects on DNA and synthetic polyelectrolytes.

---

The lack of a rigorous and thermodynamically consistent treatment of interactions between counterions and RNA has impeded a quantitative description of the self-assembly of RNA molecules (1). Although many factors contribute to the stability of a folded RNA molecule, the interplay between monovalent and divalent cations and the highly correlated nature of ion–RNA interactions make it challenging to develop an accurate and tractable theory for RNA folding thermodynamics and kinetics. The effects of monovalent ions could be accurately accounted for by using the Debye–Hückel theory (2–4). However, theoretical and computationally tractable treatments of the effects of divalent cations, such as Mg^2+^, which play an essential role in RNA structure, folding and function (5–8), have not been fully developed. Accounting for the effects of divalent ions on RNA folding requires an approach that goes beyond the use of Poisson–Boltzmann equation (9–11) in order to account for the ion size, ion–ion correlations, and the complex coordination with the phosphate groups. The simultaneous presence of monovalent and divalent ions introduces additional complexity that has to be dealt with in order to arrive at a reasonable predictive theory of RNA folding.

How ions modulate the RNA energy landscape has also been the subject of extensive experimental and theoretical studies (1, 9, 12–20). From the chemistry perspective, binding of divalent ions to the negatively charged phosphate groups could be conceptually classified into two categories: direct (or inner sphere) contact where an atom (or more) of the RNA is part of the divalent ion coordination sphere, and indirect (or outer sphere) contact where the interaction is mediated by a water molecule (5). Recent surveys of the RNA structures in the Protein Data Bank (PDB) reported the frequencies of both inner and outer spheres Mg^2+^ binding to RNA atoms (21, 22), which suggests that a theoretical model must take these interactions into account in order to describe RNA folding. However, a complete knowledge of the distribution of ions around RNA in solution is still lacking although initial insights have been provided in a recent study (23).

To arrive at an accurate model, which reliably predicts the thermodynamic properties of large RNA molecules, we first began with a sequence-dependent Three Interaction Site (TIS) coarse-grained (CG) model for nucleic acids (24), which has been adopted to study a range of problems related to RNA folding (2, 10, 25–32). Even with this simplification, the inclusion of both monovalent ions (present in excess concentration relative to divalent cations) and divalent ions explicitly is computationally demanding, although folding simulations of a 195-nucleotide *Azoarcus* ribozyme and pseudoknots have been carried out successfully (10, 33). Here, we report the folding thermodynamics of RNA molecules using simulations performed with a hybrid model in which monovalent ions are implicitly treated but divalent cations are explicitly included. To develop such a model, we resort to the Reference Interaction Site Model (RISM) theory (34–40) to obtain the potential of mean force (PMF) between divalent cations and phosphate groups. We show that this treatment is necessary to obtain accurate results for RNA thermodynamics, especially for Mg^2+^ ions, which are involved in both the inner and outer shell coordination with the negatively charged phosphate groups. Using our new model and simulations, we calculate, with high accuracy, the thermodynamics of RNA folding in the presence of divalent and monovalent cations for several RNA molecules in the folded, intermediate and unfolded states. In the process, we establish that divalent ions interact strongly with RNA molecules in the intermediate structures (41), which provides a structural interpretation of site specific interactions between RNA and ions. Our work also shows that accounting for both inner and outer sphere coordination of divalent cations with RNA is necessary to faithfully reproduce divalent cation–RNA interaction thermodynamics. The general framework, which integrates liquid state theories with molecular simulations, is applicable to investigate folding of large RNA molecules over a broad range of salt concentrations, thus vastly expanding the scope of simulations to a variety of problems in RNA biology.

## Theory

### RNA model

We adopt the Three Interaction Site (TIS) model, in which each nucleotide is modeled using three interaction sites located at the center of geometry of Phosphate (P), Sugar (S), and Base (B) (2, 3, 10, 24, 31). The energy function, *U* = *U*_BA_ + *U*_EV_ + *U*_ST_ + *U*_HB_ + *U*_EL_, takes into account the bond length and angle constraints, excluded volume interactions, secondary stacking between consecutive bases and tertiary stacking (stacking between bases that are not consecutive in the structure), both non-native and native hydrogen bond interactions. Details of the force field are given in the Supplementary Information (SI).

Previously (10), we treated all the ions (including monovalent ions) explicitly, and therefore the phosphate charge was fixed at *Q* = −*e*. Here, we employ the Debye–Hückel (DH) equation to approximate the screening effect of monovalent ions. Thus, the electrostatic interactions between the P–P and the divalent cations are written as:

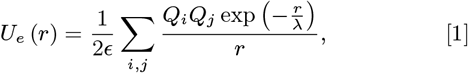

where *λ* = (8*πl*_*Bρ*1_)^*−*1/2^ is the DH screening length that depends on the number density of monovalent ions, *ρ*_1_, and 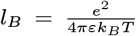 is the Bjerrum length. For divalent cations *Q*_*i*_ = +2*e*. The renormalized charge on the phosphate *Q*_P_ (*T, C*_1_, *C*_2_) depends on the concentrations of both the monovalent (*C*_1_) and divalent ions (*C*_2_), which is calculated using the counterion condensation (CIC) theory (42–44). The electrostatic interaction between the divalent ions and phosphate groups is treated precisely to account for water-mediated outer and inner shell interactions (see below). Because the X^2+^–P potential (X is Mg or Ca) includes the excluded volume interactions, we do not explicitly account for such interactions in the coarse-grained TIS force field (see the SI).

### Phosphate charge renormalization

A consequence of CIC is that the effective charge on the phosphate is reduced from −1*e*, thus softening the overall electrostatic interactions enabling the compaction and folding of the RNA. Following our earlier studies (2, 3), we include ion condensation effects for the implicitly treated monovalent ion. Since we treat divalent ions explicitly and monovalent ions implicitly, a thermodynamically consistent treatment of ion effects is needed. As the divalent ion concentration increases, the condensed divalent ions outcompete monovalent ions for the phosphate groups. This occurs because for each condensed divalent ion, approximately two monovalent ions are released, which is favored because the overall entropy of the system is increased.

In the mixed ion system, we assume that one divalent ion replaces exactly two monovalent ions, and the total RNA charge neutralized in the process is equal to those in the monovalent salt alone. In other words, if *θ*_1_ and *θ*_2_ are the number of condensed monovalent and divalent ions per phosphate group, respectively, then,

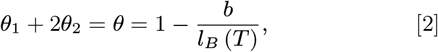

with the mean spacing between phosphate charges, *b* = 4.4 Å, a value used in our previous studies (2, 31). Although a more complicated treatment based on balance between interaction energy of ion–phosphate and entropic effect is possible (45), we find that this simple approximation works well for a broad range of ion concentrations.

A relation between *θ*_1_ and *θ*_2_ can be derived by considering the entropic cost of localizing one divalent ion *vs.* two monovalent ions. By neglecting ion–ion correlation effects, we obtain:

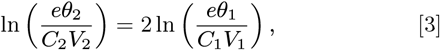

where *e* is the Euler’s number, *C*_*i*_ and *V*_*i*_ are, respectively, the bulk concentration and the effective condensation volume of ion *i*. We calculated *V*_*i*_ using:(43, 45)

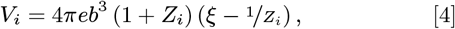

where *Z*_*i*_ is the bare charge of the ions (*Z*_1_ = +1, *Z*_2_ = +2), and the Manning parameter, 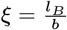. From Eqs. 2 and 3, one can determine both *θ*_1_ and *θ*_2_. Thus, the effective charge on the phosphate is *Q*_P_ (*T, C*_1_, *C*_2_) = 1 − *θ*_1_, considering only the monovalent ion condensation. We account for the contributions from the divalent cations, *θ*_2_, by treating them explicitly in the simulations. The electrostatic interactions involving the P groups are calculated using *Q*_P_(*T, C*_1_, *C*_2_), as the effective charge on the phosphate.

### Mg^2+^–P effective potential

In order to compute the Mg^2+^– P effective potential, *V*_Mg-P_ (*r*), the potential of mean force (PMF) between Mg^2+^–P derived from the RISM theory has to be modified because it is dependent on temperature and concentrations of both the monovalent and divalent ions. A number of studies have shown that it is difficult to capture the short-ranged electrostatic interactions, which has prompted others to propose several ways of separating the Coulomb potential into short and long-ranged components (46–49). The short-ranged interactions are usually determined using molecular simulations. Our approach, which is closest in spirit to a more rigorous treatment by Weeks and coworkers (50, 51), is implemented as follows. The short-ranged part of the *V*_Mg-P_ is taken to be identical to the PMF, while the long-ranged part is corrected based on the temperature and salt concentrations. We write the effective potential *V*_Mg-P_ (*r*) as,

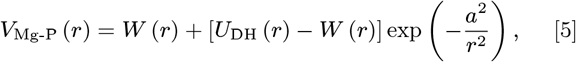

where *W*(*r*) is the PMF calculated using the RISM theory (details are in the SI), and *U*_DH_ (*r*) is the DH potential between Mg^2+^–P, accounting for the screening effect of monovalent ions. (We tried other functional forms to combine *W* and *U*_DH_, and found that Eq. 5 served our purposes in both maintaining the direct-contact interaction and smoothly merging *V* to *U*_DH_ at *r ≫ a*.) The constant, *a* = 5.0 Å, was chosen to preserve the Mg^2+^–P direct-interaction energy. Thus, at short distances a Mg^2+^ ion (or more generally, any spherical divalent cation) in proximity to the phosphate group would interact according to the PMF calculated theoretically using RISM. At large values of *r*, the Mg^2+^ ion would experience a screened phosphate charge due to the presence of monovalent ions. An advantage of our approach is that we calculate the fully equilibrated PMF between the Mg^2+^ cation and phosphate using numerical solution of the RISM equations instead of relying on MD simulations (46–49). This is particularly important for divalent ions that have slow ion-water exchange rate, and cannot be reliably implemented using conventional all-atom MD simulations (52–57).

The calculated PMF between Mg^2+^–P is shown in Fig. 1 for a solution containing 1.0 mM magnesium monophosphate. It is worth pointing out that *V*_Mg-P_ (*r*) (the red curve in Fig. 1) has two minima, one corresponding to the inner-sphere Mg^2+^ coordination to P, and the other is the outer-sphere coordination. Accounting for both the direct contact and water-mediated Mg^2+^–P interactions using Eq. 5 is necessary to calculate Mg^2+^-induced RNA folding accurately since the folding of most large RNA molecules requires both tightly bound and screening due to Mg^2+^ ions (5, 7, 10, 21). We used a similar procedure to obtain interactions for calcium *V*_Ca-P_ (*r*).

**Fig. 1.**
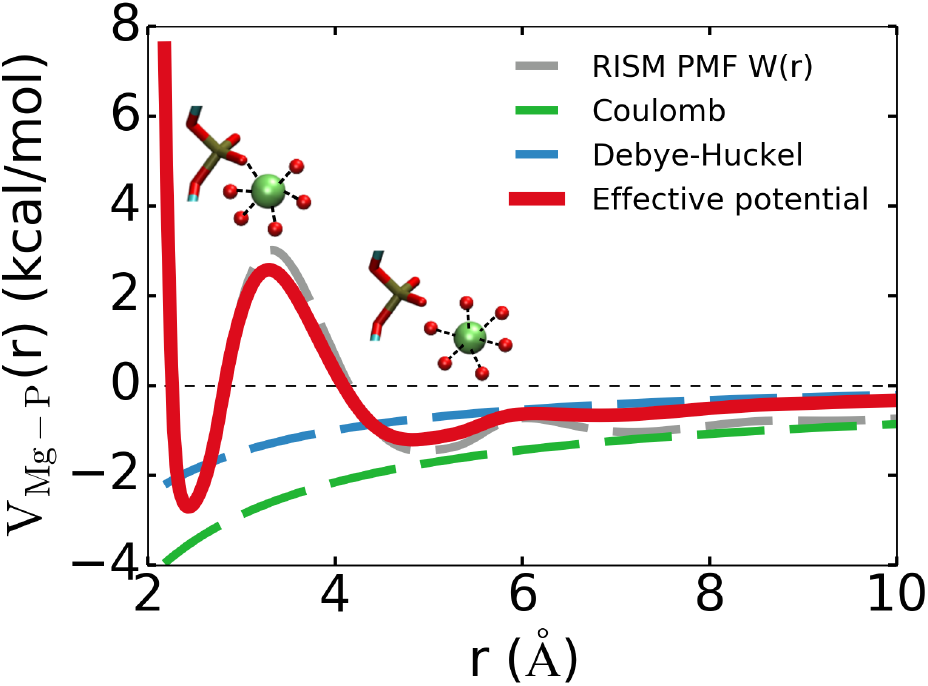
Effective (Eq. 5) Mg^2+^–P potential (red) constructed by combining the short-ranged part of the PMF (grey dashed line) with the long-ranged Debye–Hückel potential (blue dashed line). Calculations of the PMF were performed at 1.0 mM magnesium monophosphate and 25°C using the RISM theory (for details see the SI). The first minimum represents the inner shell interaction, where Mg^2+^ interacts directly with the phosphate groups. The second minimum at *r* ≈ 5 Å represents the outer shell interaction, where Mg^2+^ retains its first hydration shell.

## Results

### Determination of the parameters in the TIS RNA model

The two adjustable parameters in our RNA force field are 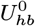 that determines the strength of hydrogen bonds, and Δ*G*_0_ that dictates the balance between stacking and hydrogen bonding (details described in the SI). Following our previous study (2), we determined the values of the two parameters by reproducing the experimental heat capacity curves of human telomerase RNA hairpin and a viral pseudoknot. The melting temperatures of the two RNA motifs are reproduced well by simulations using the new model (largest deviation is < 5*°*C) (Fig. S2). In the rest of this paper, we use this set of parameters, coupled with our treatment of divalent ion–phosphate interactions, to investigate the effects of a mixture of divalent and monovalent ions on folding of three RNA molecules. It is worth emphasizing that the same set of parameters is used for all RNA molecules over a wide range of ion concentrations.

### Preferential interaction coefficient as a function of Mg^2+^ concentration

We first performed CG simulations to probe the binding of divalent cations to the RNA, expressed in terms of the experimentally measurable ion-preferential interaction coefficient, Γ_*X*_ (where X is Mg or Ca). In the simulations, a single RNA molecule was placed in a cubic simulation box in the presence of explicitly modeled divalent cations. After equilibration, we calculated Γ_*X*_ using the Kirkwood–Buff integral (58–63),

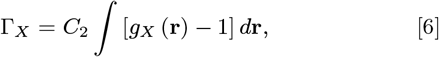

where *gx* (**r**) is the 3-dimensional distribution function of the divalent cations, reflecting the excess (or deficit) of X^2+^ relative to the bulk ion concentration, *C*_2_, in the presence of the RNA.

In Fig. 2, we show Γ_Mg_ for Beet Western Yellow Virus pseudoknot (BWYV PK), a 58-nucleotide fragment of the ribosomal RNA (rRNA), and the aptamer domain of adenine riboswitch at different concentrations of monovalent and magnesium ions. The BWYV PK folds in the presence of monovalent ions without Mg^2+^. Our simulations show that if the divalent ion concentration is increased, more of them are attracted to the PK, which is in quantitative agreement with experimental data. We also capture a more subtle experimental finding that there is a decrease of Γ_Mg_ if the monovalent ion concentration is increased from 54 mM to 79 mM, thereby effectively enhancing the competition with Mg^2+^ binding.

**Fig. 2.**
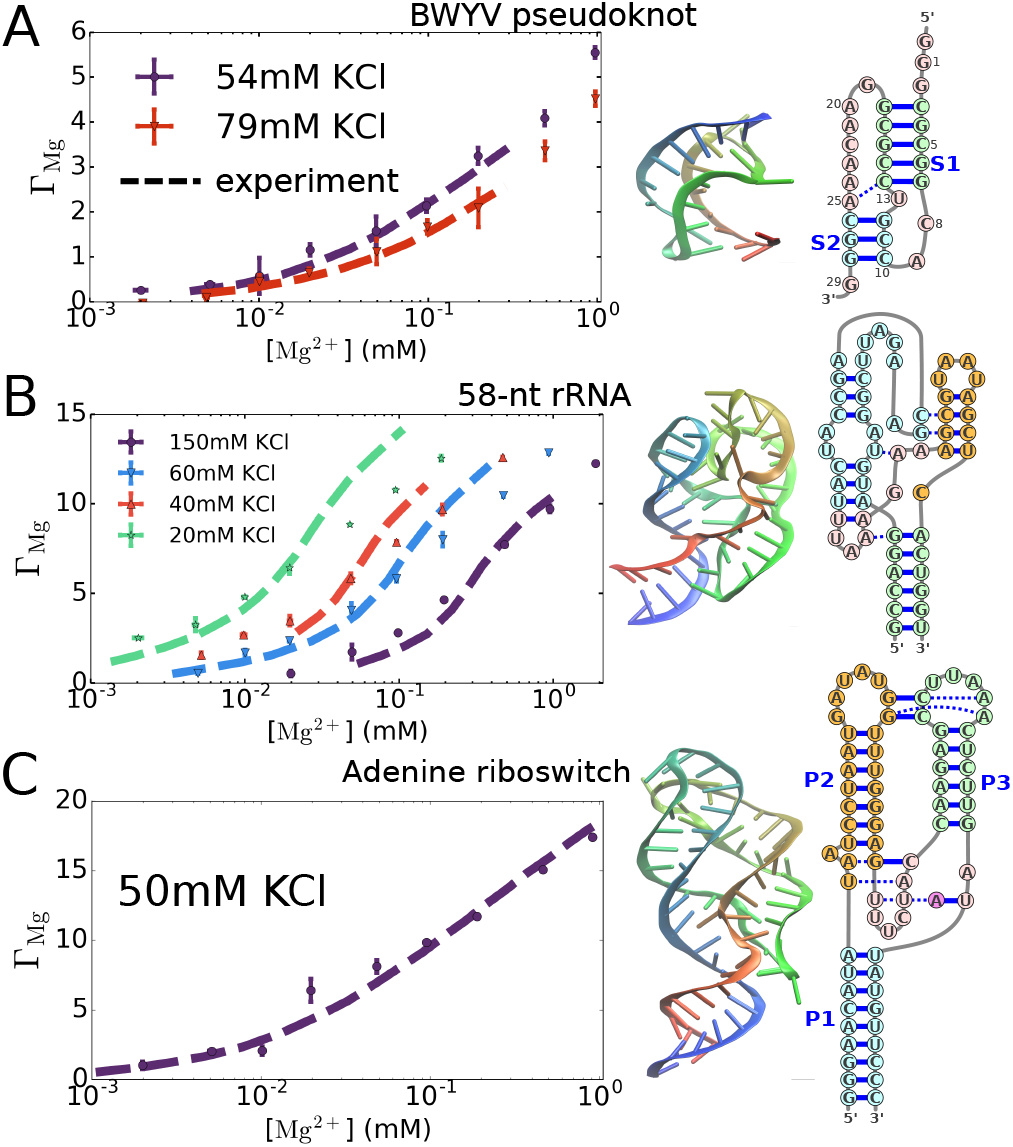
Comparison of the calculated and measured values of the preferential interaction coefficient *Γ*_Mg_ for (A) BWYV pseudoknot (64), (B) the 58-nucleotide fragment of the large subunit ribosomal RNA (65), and (C) the aptamer domain of adenine riboswitch (66) at various monovalent salt concentrations. Experimental data are taken from Refs. (67–69). The results of simulations are given by colored symbols with standard errors, and the dashed lines are data from experiments. In most cases, the error bars from the simulations are smaller than the symbol sizes. Only BWYV remains folded at all values of the concentrations of Mg^2+^. The riboswitch and rRNA partially unfold at low Mg^2+^ concentrations. At high Mg^2+^ concentrations (up to ~ 1 mM) the RNA molecules are in the folded states, whose structures are shown in the middle. Nucleotides are colored from 5’ to 3’ as red to blue. Secondary structures are shown on the right, displaying the sequences.

For the rRNA and the riboswitch, the situation is more complicated. Both of them partially unfold at low Mg^2+^ concentrations because tertiary interactions are disrupted. For these two RNA molecules, in addition to Mg^2+^ ion concentration, Γ_Mg_ also strongly depends on the state of the RNA. At low Mg^2+^ concentrations, the equilibrium shifts to extended states, which further decreases Γ_Mg_. Interestingly, our simulations quantitatively reproduce Γ_Mg_ (see Fig. 2) over a broad range of monovalent and divalent ion concentrations, for both the rRNA and riboswitch (see below for additional results for rRNA).

### Divalent ion-dependent folding free energy of BWYV pseudoknot

In a typical titration experiment, one often measures Γ_*X*_ as a function of divalent ion concentrations in the presence of excess monovalent ions (*C*_2_ ≪ *C*_1_) (41, 67, 70–73). If the RNA remains in a single state *S* (either *F*-folded, *I*-intermediate or *U*-unfolded) during the titration process, the free energy change due to the accumulation of divalent ions around RNA in state *S* is directly related to Γ_*X,S*_. For concreteness, consider the following equilibrium reaction *RNA*_*S*_ + *nX*^2+^ ⇆ *RNA*_*S*_.*nX*^2+^, showing that there is an uptake of *n* (need not be an integer) divalent cations by the RNA in the *S* state. Provided *C*_2_ ≪ *C*_1_, the free energy change associated with the equilibrium reaction given above, Δ*G*_*X,S*_, is related to Γ_*X,S*_ as,

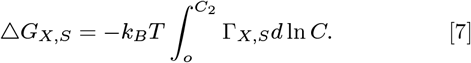

Note that Eq. 7 would not be valid if the RNA simultaneously populates different states during the titration process. For instance, if the RNA remains folded at high *C*_2_ but unfolds at low *C*_2_ (as in the rRNA and riboswitch cases), there is no obvious way to relate Γ_*X*_ and *ΔG*_*X,S*_, because Γ_*X*_ reflects the binding affinity to two (or more) states.

In order to calculate the free energy change upon folding, it is necessary to calculate *ΔG*_*X,S*_ for each state of the RNA separately, which can be done if the ensemble of RNA conformations is restricted to *S*. We choose BWYV PK for illustrative purposes because its (un)folding can be used to clearly distinguish between the folded (*F*), intermediate (*I*) and unfolded (*U*) states. Furthermore, the availability of experimental data allows us to compare directly with our simulations (67, 74).

The folded structure of the BWYV PK (Fig. 2A) has two Watson–Crick stems (*S1* and *S2*) that are connected by two small loops. The *F* state is stabilized by tertiary interactions between the two loops and the two stems. The stem *S1* has 5 G-C base pairs, while *S2* has only 3 G-C base pairs. Therefore, we anticipate that *S2* should unfold first upon increasing the temperature or lowering the salt concentration, as predicted by the stability hypothesis (75). Both experiments (74) and our previous simulations (31) have shown that BWYV does unfold by three sequential equilibrium transitions as temperature is increased, thus populating two intermediates. In one of them, there is a loss of tertiary interactions but the stems are intact. However, the probability of formation of such a state is small. Thus, for practical purposes, the overall transition to the unfolded state occurs by populating one intermediate, *F* → *I* → *U*.

In order to calculate the free energies of the folding and unfolding transitions, we first generated an ensemble of unfolded structures. We performed simulations of the *I* state by disallowing interactions between base pairs in *S2*, while preserving the full interaction for *S1*, as shown in previous studies (31, 74). The ensemble of such structures coincides with what we observed in our thermal unfolding simulations (Fig. S2). We surmise that the simulated ensemble is the one probed in the experiments (67), where all the nine 3’-terminal nucleotides were mutated to uracil, thus preventing the formation of *S2*. In the *U* state, both the stems are unfolded, which can be mimicked by disrupting all the specific interactions within the PK. This renders the PK essentially a polyelectrolyte dominated by Coulomb repulsions between the phosphate charges and secondary stackings between consecutive bases.

With the simulated ensemble of *U* and *I* structures, we calculated Γ_Mg,*I*_ and Γ_Mg,*U*_. The results are shown in Fig. 3 at two different KCl concentrations. The uptake of Mg^2+^ in the *I* and *U* states is less compared to the *F* state because they adopt more expanded conformations with spatially separated phosphate groups, thus weakening the electrostatic attraction. The calculations of Γ_Mg,*S*_ are in quantitative agreement with the experimental data. The difference ΔΓ_Mg_ = Γ_Mg,*F*_ − Γ_Mg,*I*_ between the two states is the number of Mg^2+^ uptake in the *I* → *F* transition, which is shown in the inset in Fig. 3B at two monovalent concentrations. We find a slight increase in ΔΓ_Mg_ as the Mg^2+^ concentration increases, in agreement with the direct measurement of ΔΓ_Mg_ from the fluorescence dye method (67). This is in accord with other experiments, which have shown that ΔΓ for monovalent ions also exhibits a dependence on salt concentration (44, 76). Our calculations, therefore, do not support the Wyman linkage analysis used to determine ΔΓ_Mg_, as this method yields a constant value for ΔΓ_Mg_ over the Mg^2+^ concentration range (67).

**Fig. 3.**
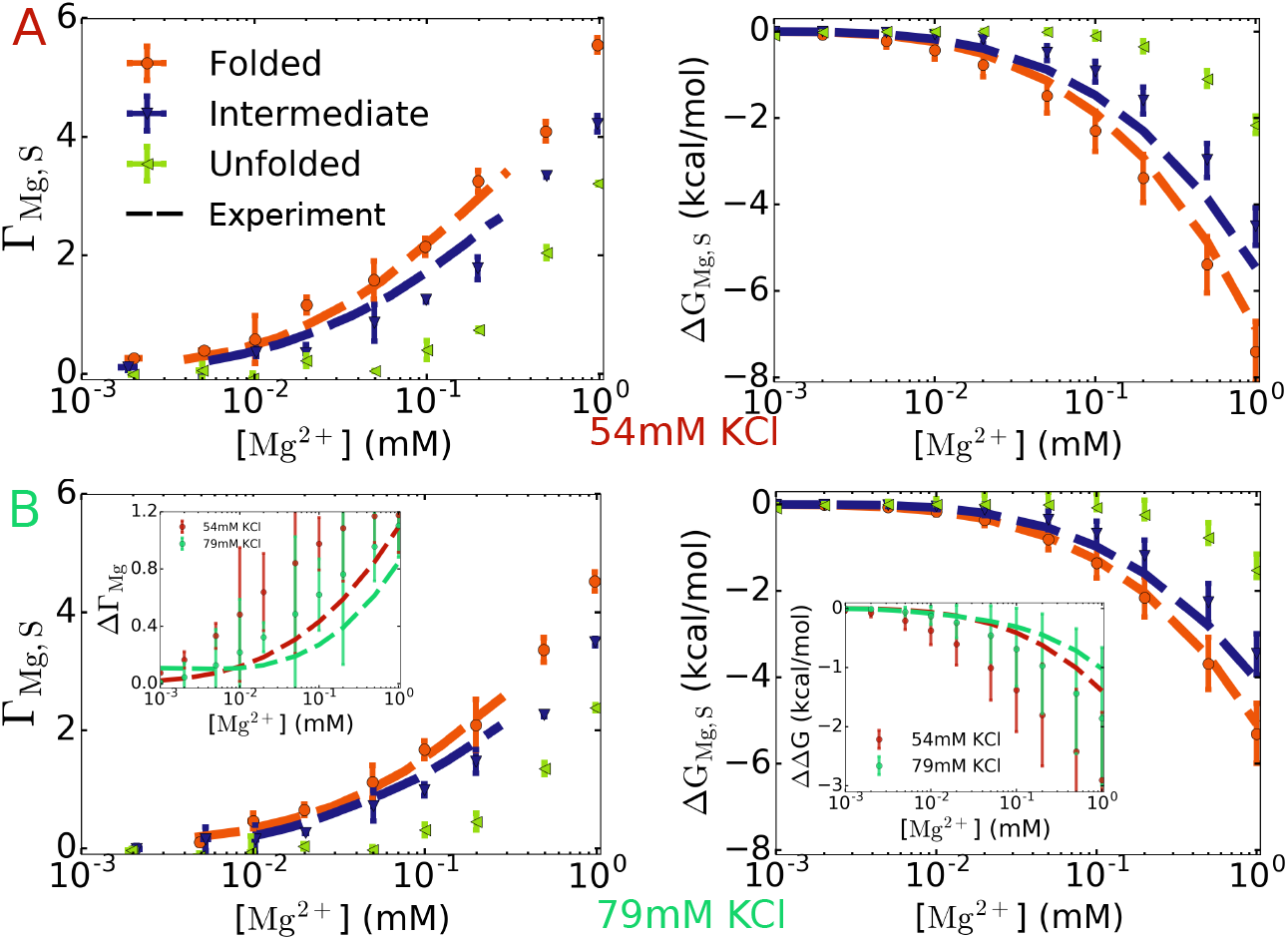
Preferential ion interaction coefficient *Γ*_Mg,*S*_ (left) and free energy of Mg^2+^–pseudoknot interaction *ΔG*_Mg,*S*_ (right) for BWYV in 54 mM (A, top) and 79 mM KCl (B, bottom), at *T* = 25°C. The calculations were performed for the folded, intermediate and unfolded states (*S* = *F*, *I*, or *U*) (see text and Fig. 4 for definition of the states). Experimental data for the *F* and *I* states are plotted as dashed lines (67). The results for the *U* state serve as a prediction of the simulations. The difference between the *F* and *I* states, *Γ*_Mg_ = Γ_Mg,*F*_ Γ_Mg,*I*_ and *ΔΔG*_F−*I*_ = *ΔG*_Mg,*F*_ − *ΔG*_Mg,*I*_, are plotted in the inset in (B). *Γ*_Mg_ (left inset) represents the number of Mg^2+^ released when BWYV transitions from the *F* to the *I* state, *ΔΔG*_F −*I*_ (right inset) is the change in the free energy of the *F* state relative to the *I* state upon addition of Mg^2+^ ions. The error bars in the inset are relatively large. However, it is clear that *Γ*_Mg_ and *ΔΔG*_F −*I*_ are not constant in the range of [Mg^2+^] here.

We then calculated the free energy changes for all the states using Eq. 7, and the results are shown in Fig. 3. Interestingly, Mg^2+^ ions are also localized near the *U* and *I* states, albeit to a lesser extent, demonstrating that it is important to characterize Mg^2+^–RNA interactions not only in the *F* state but also other relevant states along the folding pathway. For example, at 1 mM Mg^2+^ and 54 mM KCl, *ΔG*_Mg,*U*_ ~ −2 kcal/mol, *ΔG*_Mg,*I*_ ~ −4 kcal/mol while *ΔG*_Mg,*F*_ ~ −7 kcal/mol. One way to quantify the effect of Mg^2+^ addition on the folding process is to compute *ΔΔG*. For instance, for the *F* → *I* transition, *ΔΔG*_*F −I*_ = *ΔG*_Mg,*I*_ − *ΔG*_Mg,*F*_. The stabilization of the *F* state relative to the *I* state or *U* state caused by Mg^2+^ addition (*ΔΔG*_*F −I*_ or *ΔΔG*_*F−U*_) is therefore ~ −3 and −5 kcal/mol, respectively. The relatively small value of the Mg^2+^ dependence on the folding free energy is likely due to the small size of this PK, whose folded state is stable even in the absence of Mg^2+^ (see Fig. S2B and S5).

### Free energy of BWYV pseudoknot using thermodynamic cycle

In Fig. 4, we illustrate how the free energy data shown in Fig. 3 in conjunction with the RNA folding free energy data obtained by varying the temperature could be used to construct a folding free energy diagram for BWYV at 0.2 mM Mg^2+^. Similar diagrams at arbitrary concentration of Mg^2+^ can be generated. The vertical free energy differences (Fig. 4) are from WHAM analysis of multiple-temperature simulations with or without Mg^2+^ (see SI for additional details). The horizontal free energy differences are Mg^2+^–RNA free energies as in Fig. 3. The Mg^2+^-dependent free energy of stabilization is evaluated by two ways using the thermodynamic cycle,

**Fig. 4.**
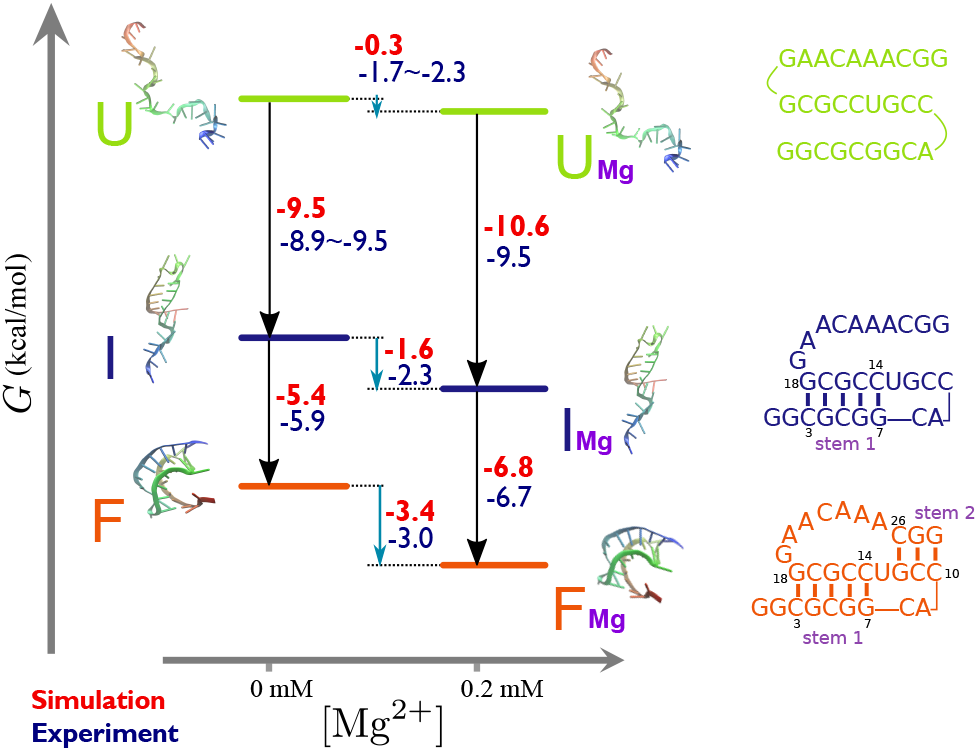
Folding free energy (in kcal/mol, 25°C) diagram of BWYV pseudoknot at 54 mM KCl in the absence (left) or presence (right) of 0.2 mM Mg^2+^. Theoretical values are in red and experimental values are in blue. Experimental data for. 6. *ΔG*_U-I_ (no Mg^2+^) and *ΔG*_Mg,*U*_ are not available. For *ΔG*_U-I_, data for 40 mM and 74 mM KCl are reported with the hope that they should bracket the 54 mM KCl data. *ΔG*_Mg,*U*_ for experiment is then evaluated based on the other free energies in the cycle. Folding free energies (black arrows) are calculated from thermal denaturation simulations. Mg^2+^–RNA free energies (blue arrows) are taken for each state at 0.2 mM Mg^2+^ from Fig. 3.

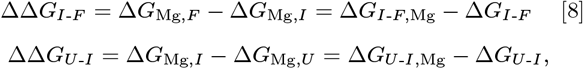

where *ΔG*_*α-β*,Mg_ and *ΔG*_*α-β*_ are, respectively, the free energy difference between the *α* and *β* states (*α, β* is *F*, *I*, or *U*) in the presence and absence of Mg^2+^. The calculated free energies are in remarkable agreement with experiments and are consistent with each other (*ΔΔG* values estimated by two different methods give similar values, with the errors ±0.4 kcal/mol). Given that they are independently determined, it shows that our theory can be reliably used to study thermodynamics of Mg^2+^-induced RNA folding.

### Mg^2+^-induced folding of the 58-nucleotide rRNA

We investigated the folding of the 58-nt rRNA, which, although folds at high (~ 1.6 M) monovalent ion concentrations, requires Mg^2+^ (68). We carried out simulations of the 58-nt rRNA as a function of different combinations of monovalent and divalent ion concentrations. Fig. 2B already shows that we can quantitatively account for the dependence of Γ_Mg_ as a function of both monovalent and divalent ion concentrations. To fully characterize the folding of the rRNA fragment, we calculated the radius of gyration, *R*_*g*_, as a function of Mg^2+^ concentration at four concentrations of KCl. Fig. 5A shows that *R*_*g*_ values decrease continuously from ~ 25 Å at [Mg^2+^] = 2 *µ*M to ~ 16 Å at high Mg^2+^ concentrations, which is consistent with experimental measurements at 40 mM KCl (shown as red bars in Fig. 5) (68). Interestingly, our calculations show that the concentration of KCl has minimal effect on the dependence of *R*_*g*_ on Mg^2+^, even at low Mg^2+^ concentrations at which the RNA is partially unfolded. This is because at these KCl concentrations, the secondary structure of the rRNA is fully formed (Fig. 5B). Therefore, adjusting the monovalent ion concentration does not considerably assist folding since it requires Mg^2+^ for tertiary structure formation. Fig. 5B shows the average fraction of native contacts for the rRNA, *Q*, as a function of Mg^2+^ concentrations. In accord with *R*_*g*_ analysis, the *Q* values at low [Mg^2+^] are small, fluctuating around ~ 0.35. At these Mg^2+^ concentrations, the secondary structure of the rRNA is completely stabilized without forming tertiary interactions. At high [Mg^2+^], *Q* increases and reaches ~ 0.7. Just as found for *R*_*g*_, we also observe a negligible dependence of *Q* on monovalent ion concentration. Thus, both the order parameters (*R*_*g*_ and *Q*) show that rRNA formation, which does not depend much on KCl concentrations, is only moderately cooperative.

**Fig. 5.**
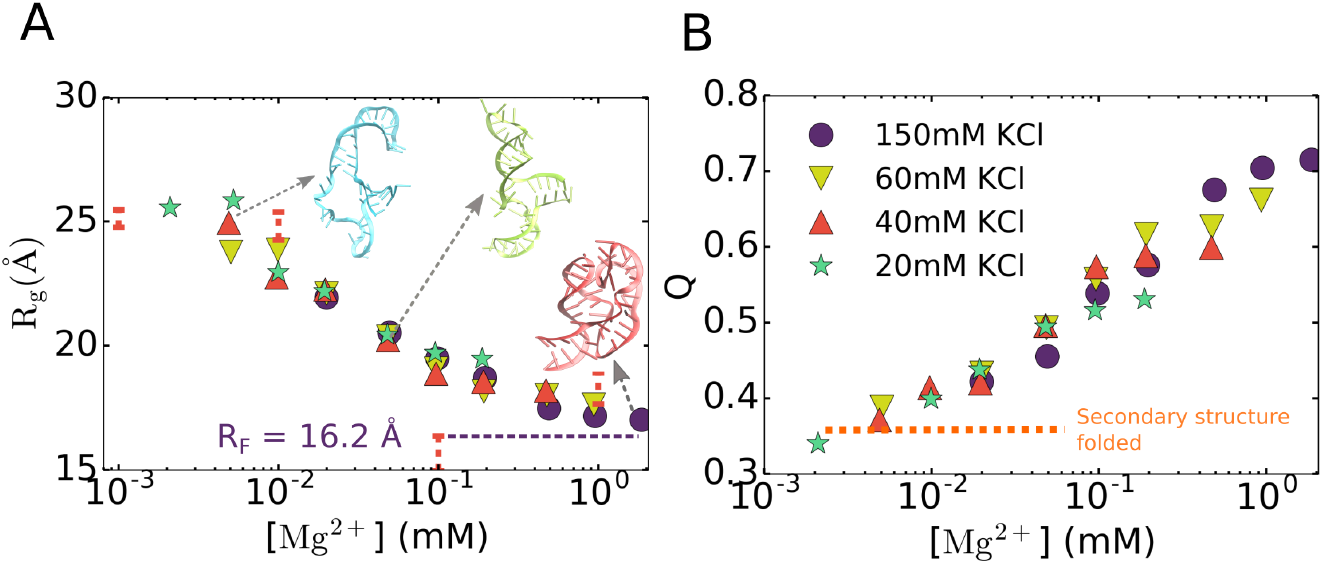
Mg^2+^-induced folding of the 58-nt fragment of ribosomal RNA. (A) Compaction of rRNA as the Mg^2+^ concentration increases. The colors correspond to four monovalent ion concentrations (shown in B). *R*_*F*_ is the value of *R*_*g*_ for the *F* state, calculated from the PDB structure (PDB ID 1HC8). Experimental measurements of *R*_*g*_ in 40 mM KCl are plotted as vertical red error bars (68). Some representative structures from the simulations are also shown. (B) Average fraction of native contacts vs. Mg^2+^ concentration. The horizontal dashed line at around *Q ≈* 0.35 indicates complete secondary structure formation with no tertiary interactions.

### Free energy changes upon folding

In order to obtain the folding thermodynamics of rRNA we calculated *ΔG*_Mg,*S*_ for each state of the 58-nt rRNA by assuming that the folding transition occurs sequentially, *U* → *I* → *F*, where the *I* state is comprised only of secondary structure of three stems connected by small loops (see Fig. 6). It is possible that for such a complex RNA, more than one intermediate state could be populated during the (un)folding process. However, we chose only one intermediate state to separate the effect of divalent cations on secondary and tertiary interaction formation. The transition *U* → *I* only involves secondary structure formation, while *I* → *F* requires the formation of only tertiary interactions. As before, we performed simulations for each state by constraining the ensemble of RNA structures in such state. We emphasize that these simulations are completely different from the simulations above (from which we calculated Γ_Mg_, *R*_*g*_ and *Q*), which were performed using all the sampled conformations. The constrained simulations were only used to calculate Γ_Mg,*S*_, Mg^2+^–RNA free energies, *ΔG*_Mg,*S*_ and *ΔΔG*_*α-β*_ (where *S*, *α*, and *β* is *F*, *I* or *U*).

**Fig. 6.**
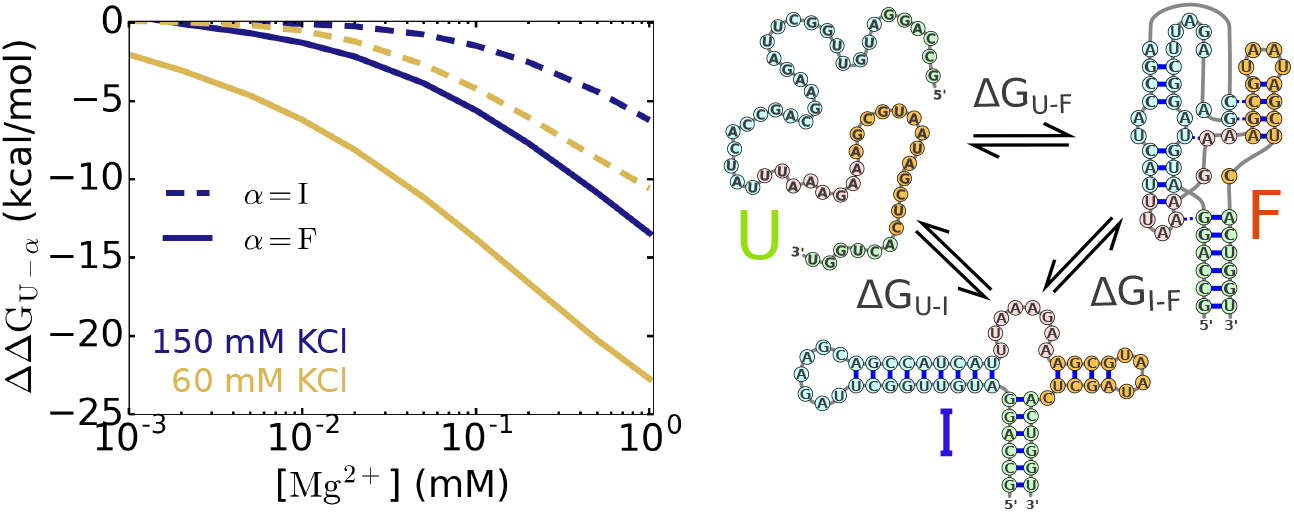
Stabilization free energies, *ΔΔG* (defined in Eq. 8), of the intermediate state (dashed lines) or the folded state (solid lines) relative to the unfolded state upon addition of Mg^2+^ ions for rRNA. *ΔΔG*_*I-F*_ is the difference between these two free energies *ΔΔG*_*I-F*_ = *ΔΔG*_*U-F*_ − *ΔΔG*_*U-I*_. In the *I* state, secondary structures are formed. Therefore, *ΔΔG*_*U-I*_ and *ΔΔG*_*I-F*_ are, respectively, are the stabilization free energies of Mg^2+^ on secondary and tertiary structure formation.

In Fig. 6, we show the results for the stabilization free energies of the folding transitions upon addition of Mg^2+^ ions, *ΔΔG*_*α-β*_, calculated using Eq. 8. Data for Γ_Mg,*S*_ and *ΔG*_Mg,*S*_, which were used to compute *ΔΔG*_*α-β*_, can be found in Fig. S7. *ΔΔG*_*α-β*_ shows the change of the relative stability of the two states *α* and *β* upon addition of Mg^2+^ on the *α ↔ β* transition. We explicitly show only *ΔΔG*_*U-I*_ and *ΔΔG*_*U-F*_ curves, but *ΔΔG*_*I-F*_ can be calculated as the difference between these two curves, as *ΔΔG*_*I-F*_ = *ΔΔG*_*U-F*_ − *ΔΔG*_*U-I*_. It is obvious that the higher the [Mg^2+^] is, larger is the effect of Mg^2+^ ions on all three transitions since all *ΔΔG* values decrease (increase in magnitude) as [Mg^2+^] raises. Therefore, higher [Mg^2+^] induces a shift in the equilibrium towards more compact states (*U* → *I* → *F*). The magnitude of *ΔΔG* for rRNA is quite large compared to BWYV, indicating the dramatic dependence of the rRNA folding on Mg^2+^. For rRNA in 60 mM KCl, adding 0.1 mM Mg^2+^ leads to *ΔΔG*_*I-F*_ ≈ −8 kcal/mol and *ΔΔG*_*U-F*_ ≈ −13 kcal/mol. In comparison, for BWYV at 54 mM KCl, those values are *ΔΔG*_*I-F*_ ≈ −1.4 kcal/mol and *ΔΔG*_*U-F*_ ≈ −2.2 kcal/mol, respectively. On the other hand, if one instead increases the concentration of monovalent ions, the values of *ΔΔG* become smaller (Fig. 6), which also happens in the BWYV PK (shown in the inset of Fig. 3B). At 150 mM KCl, adding 0.1 mM Mg^2+^ into the solution of rRNA only leads to *ΔΔG*_*I-F*_ ≈ −4 kcal/mol and *ΔΔG*_*U-F*_ ≈ −6 kcal/mol, respectively.

### Comparison between Mg^2+^ and Ca_2+_ ions

We also studied the effect of the cation size (Mg^2+^ *vs.* Ca^2+^) on RNA folding. We computed the Ca^2+^–P effective potential, *V*_Ca-P_ (*r*), using the same procedure used to obtain *V*_Mg-P_ (*r*). Fig. 7A compares the effective potentials of the two ions. A major difference between *V*_Ca-P_ (*r*) and *V*_Mg-P_ (*r*) is that the barrier separating the inner shell and outer shell binding in Ca^2+^ is much lower than in Mg^2+^. This difference arises because the charge density of Mg^2+^ is much higher than Ca^2+^, which results in orders of magnitude difference in the water exchange kinetics between Mg^2+^ and Ca^2+^ (77). It is also in accord with the observation that the interaction of Mg^2+^ with water in the first hydration shell is stronger than in Ca^2+^ (Fig. S10).

**Fig. 7.**
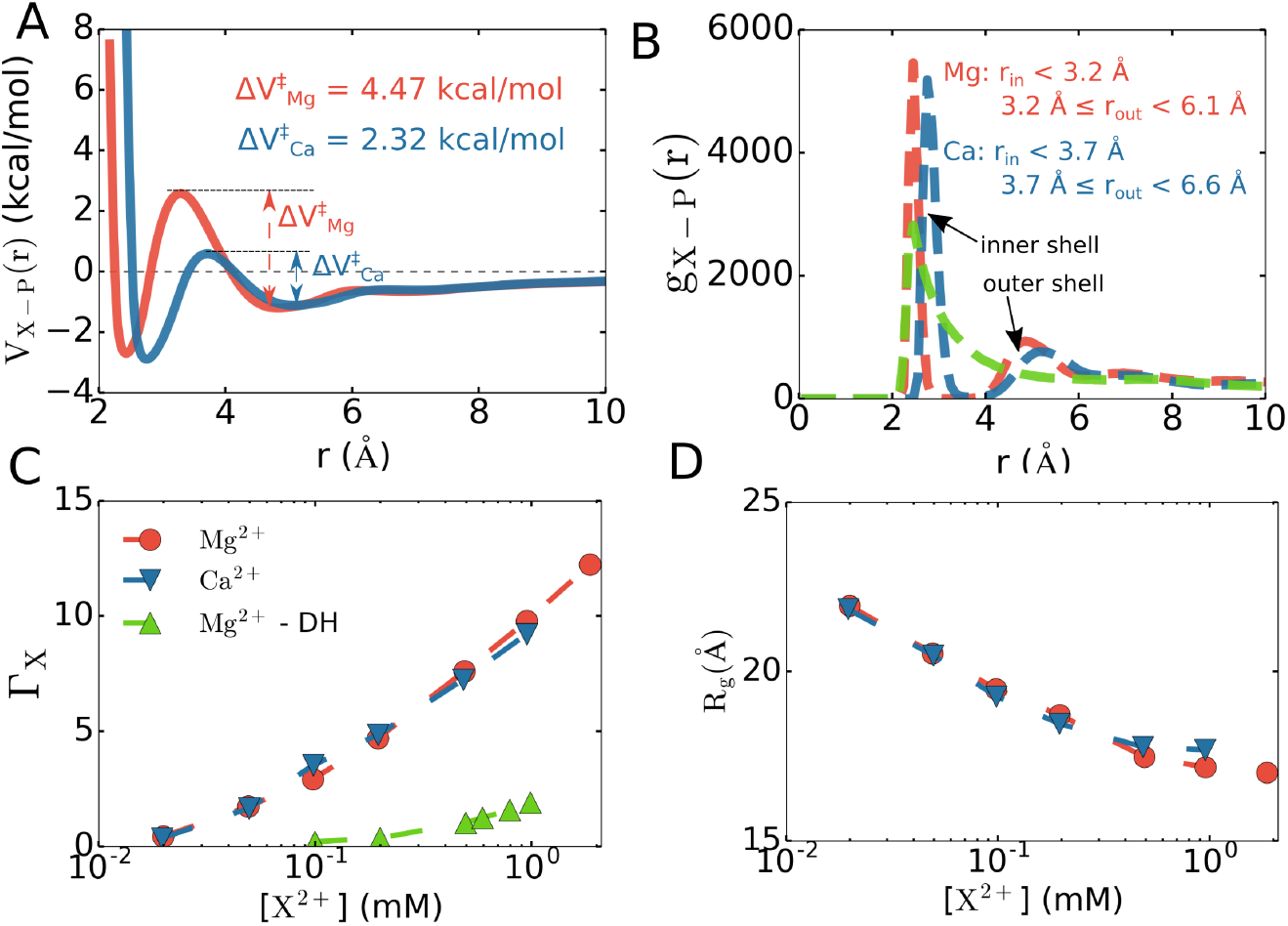
Comparison between Mg^2+^ and Ca^2+^ binding to phosphate groups and RNA. (A) Effective potentials for Mg^2+^ and Ca^2+^ with the phosphate group show that the transition barrier between inner shell-outer shell coordination for Ca^2+^ is considerably lower than for Mg^2+^. However, the depth of the inner and outer shell minima are comparable. (B) Radial distribution function between the ions and phosphate groups in BWYV. Also shown the results for Mg^2+^ ions in which Mg^2+^–phosphate interactions are modeled using the Debye–Hückel potential (green). (C) Preferential interaction coefficient Γ_*X*_ and (D) Radius of gyration *R*_*g*_ computed for 58-nt rRNA. Only data for 150 mM KCl is presented here, data for other KCl concentrations can be found in the SI (Fig. S6). The rRNA reaches its native states at high divalent ion concentrations using our model, but does not easily fold at any Mg^2+^ concentration should the Debye–Hückel potential be used.

Fig. 7B shows the radial distribution function, *g*_X-P_ (*r*), between the divalent ions and phosphate groups in BWYV PK. It is obvious that our model accounts for both types of binding. The presence of inner and outer shell binding of Mg^2+^ to P is indicated by two visible peaks in *g*_Mg-P_ (*r*). The first peak is located at *r* ~ 2.4 Å and the second is at *r* ~ 4.8 Å. The slow decay of *g*_Mg-P_ (*r*) towards 1 is due to the presence of other phosphates in the RNA. The peaks for Ca^2+^ are very similar, and are shifted slightly upwards towards longer distance (2.7 Å and 5.1 Å), indicating that Ca^2+^ has a comparable affinity for the phosphate groups.

We also investigated the folding of the rRNA in the presence of Ca^2+^ ions. Due to the similar affinity of the two ions towards phosphate groups, there is little difference in the folding behavior of rRNA between Ca^2+^ and Mg^2+^ in terms of ion accumulation, fraction of native contacts and global size (Fig. 7C and D). It is possible that the difference between the two ions is only apparent in the case of more complex RNAs, such as group I intron ribozyme (10), where the folded state is highly compact and there is not sufficient room in the core of the RNA to accommodate larger ions, and therefore replacing Mg^2+^ by Ca^2+^ in these cases would destabilize the folded state. Nonetheless, we find an interesting difference in the nature of binding of the two ions: Ca^2+^ dehydrates readily due to its lower charge density, and binds the phosphate groups directly in the inner shell, while Mg^2+^ coordination, with a higher charge density, is roughly similar between the inner and outer shell. A more detailed study will be reported in a subsequent publication.

In addition, we also show in Fig. 7 data for Mg^2+^ assuming that the Mg^2+^–P interaction is given by the DH potential (green curves). In the *g*_Mg-P_ (*r*) plot (Fig. 7B), the DH potential completely misses the second peak and the first peak is also much lower compared to our model. This leads to lower affinity with the phosphate groups, resulting to fewer ions accumulated around the RNA. When applying the DH potential to rRNA folding, we find that the RNA does not fold, but rather adopts much more extended conformations, which is directly related to the small uptake of ions, at all Mg^2+^ concentrations (Fig. 7C and D). Our model thus reveals the importance of treating Mg^2+^–P interaction accurately in order to faithfully capture both structural and thermodynamic features of Mg^2+^-assisted RNA folding.

### Importance of outer-sphere Mg^2+^-P coordination

One of the key predictions of this work is that accurate predictions of RNA folding thermodynamics require a consistent description of both the inner and outer sphere coordination of divalent cations to phosphate groups. In order to assess the importance of the outer sphere coordination, we created a potential that retains the inner shell interaction between Mg^2+^–P, while smoothly joining the outer shell interaction with the DH potential (see Fig. 8A). In so doing all the outer shell interactions between Mg^2+^–P are eliminated. The barrier between inner sphere-outer sphere in the modified potential is lower than in the original potential. We believe that the smaller barrier would only alter the kinetics of water and ion exchange around Mg^2+^, and should not significantly affect the folding thermodynamics quantities. The outer-sphere coordination interaction is only weakened ~0.2 kcal/mol in the modified potential.

**Fig. 8.**
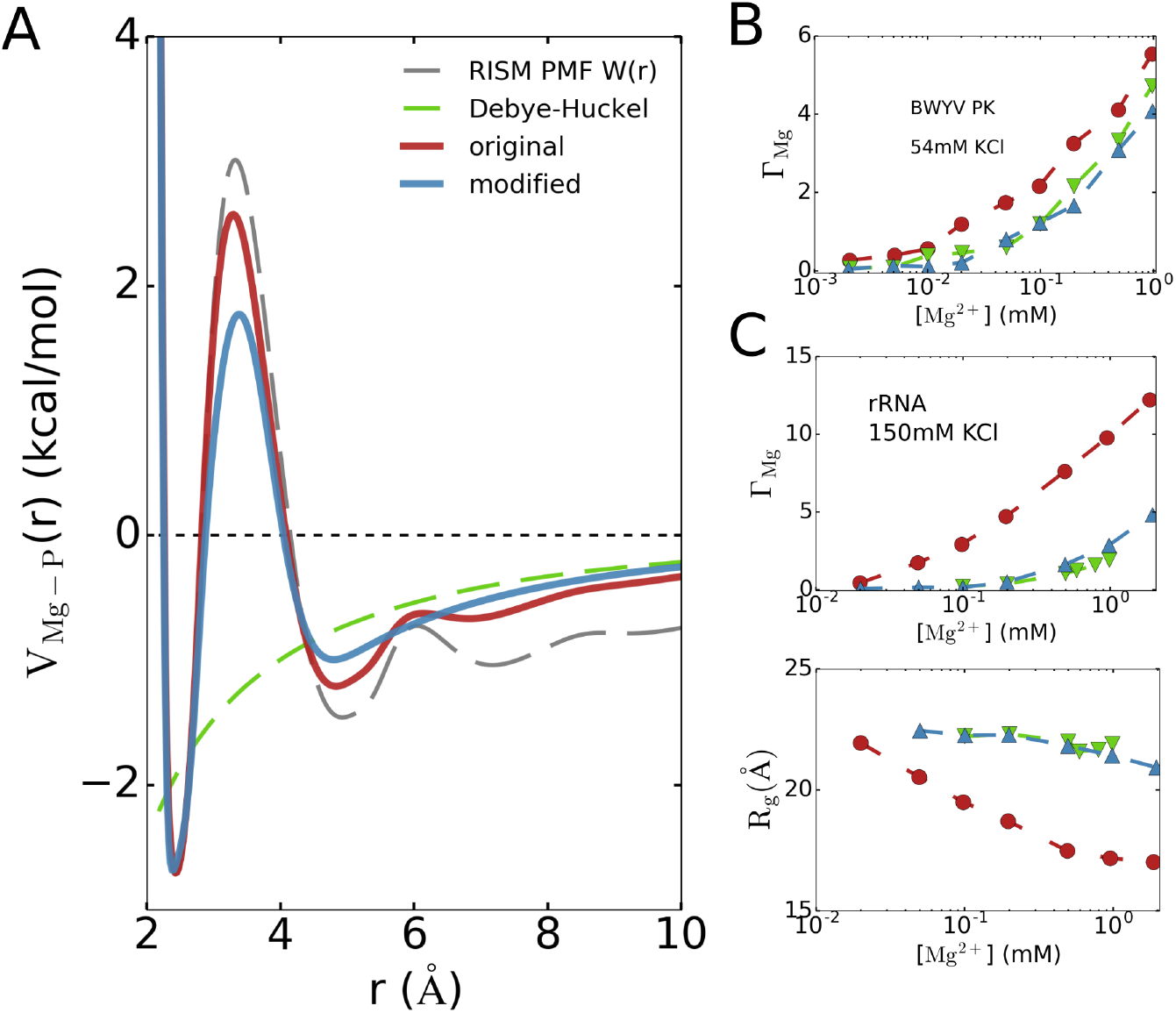
Importance of treating both inner- and outer-sphere coordination. (A) Modified effective potential for Mg^2+^ with softened outer-sphere coordination interaction, while keeping the inner-sphere coordination interaction intact. Comparison of (B) Preferential interaction coefficient Γ_Mg_ for BWYV PK at 54 mM KCl and (C) Γ_Mg_ and *R*_g_ for 58-nt rRNA at 150 mM KCl using the modified potential. Despite a tiny difference in the outer-sphere coordination (~0.2 kcal/mol), the values of Γ_Mg_ decrease significantly from the original model. In rRNA, the modified potential even cannot fold the rRNA at relevant Mg^2+^ concentrations, as can be seen in the high values of *R*_*g*_.

The simulations using the modified potential reveal a large impact on Mg^2+^–RNA interactions and RNA folding. The substantial changes are illustrated in Fig. 8B, comparing the values of Γ_Mg_ for BWYV in 54 mM KCl. A seemingly small softening of the outer-sphere coordination interaction leads to a significant decrease in Γ_Mg_. This shows, rather vividly, that the Mg^2+^–RNA interaction, and therefore the Mg^2+^–RNA free energy, is extremely sensitive to the Mg^2+^–P interaction. In rRNA, the situation is even more pronounced (Fig. 8C). Because the folding of the rRNA depends dramatically on Mg^2+^, weakening the outer-sphere coordination interaction modestly prevents the folding of rRNA even at high [Mg^2+^] due to insufficient number of condensed Mg^2+^. Interestingly, the use of the modified potential results in insignificant compaction of rRNA even at the highest Mg^2+^ concentration (lower panel in Fig. 8C). Surprisingly, there is negligible difference the predictions for *R*_*g*_ between the predictions using the DH potential and the modified potential. The results in Fig. 8B and Fig. 8C show that accurate predictions for RNA folding thermodynamics requires accounting for both inner and outer sphere coordination of Mg^2+^ with phosphates.

## Discussions

In this study, we have introduced a new method to capture the impact of a solution containing a mixture of monovalent and divalent cations on RNA folding. In our model, based in part on liquid state theory, divalent cations are treated explicitly while the screening effect of monovalent salt is treated implicitly. We reasoned that since the DH theory works well at long range (in fact, the DH theory gives asymptotically correct results at long distances), we could improve the divalent cation interactions by modifying the hard to describe short-ranged interactions. To obtain the short-ranged interactions, we used RISM theory to compute the PMF between divalent cation–phosphate group and combined it with the long-ranged part of the DH potential to obtain the effective potential *V*_X-P_ (*r*). Applications to three RNA molecules, with different sequences and structures, illustrate that our theory quantitatively reproduce the Mg^2+^ preferential interaction coefficients and Mg^2+^–RNA free energies for different combinations of ion concentrations. The transition free energies as the RNA traverses along the folding pathway are also in remarkable agreement with experimental data. The simulations not only reproduce thermodynamics of divalent cation binding to the RNA but also recapitulate the correlation between divalent cation concentration and RNA folding. In addition, we presented the effects of divalent cations on secondary and tertiary structure formations for an RNA construct and its dependence on monovalent concentrations.

The difference in the divalent cation–RNA interactions between our model and DH theory occurs only in the short-ranged part, *r*_X-P_ ≤ 10 Å (little difference exists from *r* ~ 6 − 10 Å in Fig. 1). Nonetheless, such a difference proves to be very crucial in obtaining the correct divalent cation binding free energies, as it translates to at least ~ 2.0 kcal/mol deviation of *ΔG*_Mg,*F*_ for BWYV (Fig. S9). We also show that in order to obtain the correct divalent ion–RNA binding free energies, it is important to take into account both inner and outer shell interaction accurately. A deviation of only 0.2 kcal/mol in the divalent cation–P potential could lead to a substantial decrease in the number of bound ions. In complex RNAs whose folding depends on divalent ions, loss of bound ions could even cause the RNAs to be thermodynamically unstable even at elevated ion concentrations. It is also worth stating that since our model treats monovalent ions implicitly, it has a large advantage over fully explicit ion models in terms of simulation performance, and could be used to study much more larger RNA molecules including RNA-protein interactions. Indeed, our theory is sufficiently general that it can be applied to calculate ion (with arbitrary valence and size) effects on DNA as well as synthetic polyelectrolytes and polyampholytes.

Although no other existing computational model can be used to calculate ion-dependent folding thermodynamic properties of RNA of arbitrary size and sequence accurately as we have done here, our theory is not without limitations. For example, we considered only the divalent ion–phosphate interaction and neglected interactions with the bases, which might be important as the size of RNA molecules increase. It is suspected that Mg^2+^ ion interacts with electronegative atoms in the base moiety (both inner shell and outer shell) (21, 22). However, it is unknown if such interactions are relevant for RNA folding thermodynamics. In addition, our RNA force field includes non-native interactions only in a limited manner. Nevertheless, the remarkable agreement between the theoretical predictions and experiments opens new ways to quantitatively probe ion-induced folding of RNAs regardless of their sizes and sequences.

## Conclusions

We have proposed a new theory of divalent ion–phosphate interactions, based on concepts in liquid state physics, for use in coarse-grained simulations of RNA folding in the presence of explicit divalent cations while the screening effect of monovalent salt is treated implicitly. Because our model accounts for both the inner and outer sphere coordination of divalent cations with the RNAs using RISM, the theory quantitatively reproduces divalent cation-dependent free energies for folding transitions and the correlation between the divalent cation binding and RNA folding. The success of our general method, which integrates liquid state theories and coarse-grained TIS model for RNA, is widely applicable to a variety of problems in RNA biology in which divalent cations play an important role. Finally, the theory could also be used to treat the effects of spherical and non-spherical ions on the conformations of RNA as well as DNA.

## Materials and Methods

Full details of RISM theory, the calculation of divalent cation–phosphate potential, RNA coarse-grained force field and simulation details are provided in the SI.

## ACKNOWLEDGMENTS

We are grateful to Tom Record for insightful discussions. HTN thanks Natalia Denesyuk for providing the source code for the previous model, Mauro Mugnai and Debayan Chakraborty for several fruitful discussions. This work was supported by a grant from the National Science Foundation (CHE 19-00093) and the Welch Foundation (F-0019) through the Collie– Welch chair. We are thankful to the Texas Advanced Computing Center (TACC) for providing computational resources.

## Supplementary Information

### Supporting Information Text

The Supplementary Information is divided into the following sections. The details of the theory used to numerically solve the RISM integral equation, needed to calculate the potential of mean force between divalent ions and the phosphate groups are given in Section 1. Section 2 describes the development of the RNA force field, which treats divalent cations explicitly and the monovalent ions implicitly. We also describe the methods used to determine the values of the force field parameters. The simulation details and methods used to analyze the data are given in Section 3. Additional tests of the validity of the theory-based construction of the RNA model and simulations are contained in Section 4.

#### 1. Divalent ion–phosphate potential of mean force

##### Reference Interaction Site Model (RISM)

Accurate simulations using coarse-grained models of even modestly sized RNA molecules in explicit monovalent and divalent cations is computationally demanding (1). In order to simplify the problem, while still retaining high level of accuracy achieved previously (1), we treat the electrostatic effects due to monovalent ions implicitly. This leaves us with the task of calculating the effective interactions between the divalent cations and RNA. Our primary goal is to calculate the potential of mean force (PMF), *W* (*r*), between Mg^2+^/Ca^2+^ and phosphate (Eq. 5 in the main text), which can be used in simulations of divalent cation-induced folding of RNA. In order to calculate *W* (*r*), we resort to the well-known RISM theory, which was developed to calculate the equilibrium site-site distributions of polyatomic liquids and their associated thermodynamic properties (2–8). The theory has two versions: one is a 1-dimensional RISM (or 1D-RISM) and the other is a 3-dimensional RISM (3D-RISM). The former provides the radial distribution functions, *g*_*ij*_ (*r*), between every interaction site in the system. The latter couples the 1D radial information and the 3D structure of the biomolecule to yield the solvent structure around the biomolecule in the form of a 3D site distribution function, *g*_*i*_ (*r*), for each solvent site. Because the theory and implementation are widely known, here we only give a very brief summary of the 1D-RISM that is most directly relevant to our work.

We begin with the Ornstein–Zernike (OZ) equation:

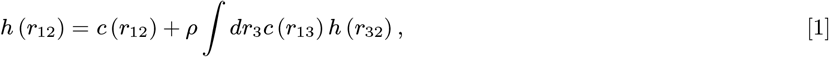

where *r*_*ij*_ is the distance between particles *i* and *j*, *c* is the direct correlation function, and *h*–the total correlation function–is related to the pair distribution function, *h*_*ij*_ (*r*_*ij*_) ≡ *g*_*ij*_ (*r*_*ij*_) − 1. In order to solve the OZ equation, it is necessary to use an appropriate closure relation connecting *h* and *c*, which we write as:

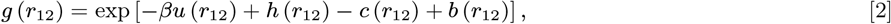

or in a short form *g* = exp [−*βu* + *h − c* + *b*]. In the above equation, *u* is the potential energy function, 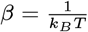 (*k*_*B*_ is the Boltzmann constant and T is the temperature), and *b* is an unknown “bridge function”. In the hypernetted-chain approximation (HNC), *b* is zero, giving:

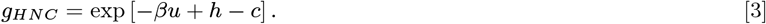

The HNC closure gives good results for ionic and polar systems, but not for neutral systems. Moreover, it is difficult to find converged solutions (6, 9, 10). To resolve these problems, Kovalenko–Hirata (KH) introduced the following closure relation: (11)

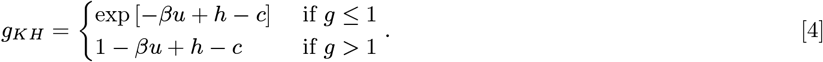

The partial series expansion of order-n (PSE-n) offers a way to interpolate between Eqs. 3 and 4, which improves the results of the KH closure while circumventing the convergence issues associated with the HNC closure: (12)

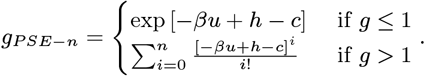

Hence, KH is the special case of PSE closure when *n* = 1. In the limit *n* → ∞, HNC is obtained.

In RISM (as implemented in the Amber force field), the standard Coulomb and Lennard–Jones interaction are used for the pair-wise non-bonded potential: (13)

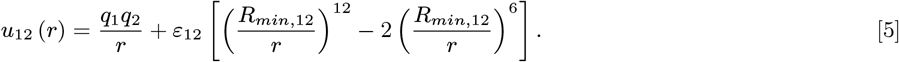

Recently, a 12-6-4 LJ potential was proposed to account for the ion-induced dipole moment interaction, which proved to be important for divalent ions (14). With this scheme, the LJ interaction (the second term in Eq. 5) is modified as:

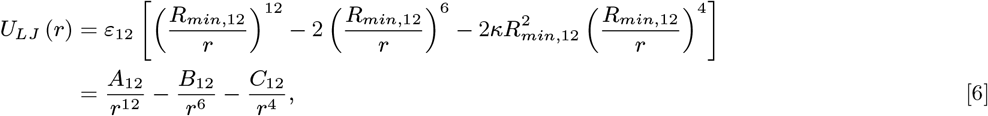

where the attractive term 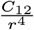 approximately accounts for the charge-induced dipole interaction. For highly charged systems, the potential in Eq. 6 yields accurate values of hydration free energies, ion-oxygen distances in the first hydration shell, and coordination numbers for divalent ions (and later, extending to trivalent and tetravalent ions) (15). A modified parameter set for Mg^2+^ was subsequently developed to balance the interaction between Mg^2+^ and water, Mg^2+^ and specific sites on nucleic acids (16). Here, we adopt these modifications in RISM and treat only the interaction involving Mg^2+^ (namely, Mg^2+^–water and Mg^2+^–P) using the 12-6-4 potential (Eq. 6), while the rest (water–water, water–P, P–P and Mg^2+^–Mg^2+^) are modeled using the standard 12-6 potential, 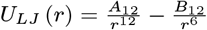. (The small polarizability of Mg^2+^ allows us to neglect Mg^2+^–Mg^2+^ interactions.) A similar procedure is used to calculate Ca^2+^ –P interactions.

The RISM equations (Eqs. 1 and 2) are solved iteratively until converged results are obtained. We are interested only in the probability distribution between Mg^2+^ and phosphate, from which the PMF is computed using *W* (*r*) = −*k*_*B*_*T* ln *g* (*r*).

##### Numerical solution of the RISM calculation

The PMFs between the divalent cations and phosphates were calculated using the 1D-RISM implemented in Amber (13) by modifying the `rism1d` code to include the potential in Eq. 6. The theory requires bulk concentrations and topologies (bond lengths and bond angles) of every molecule and ion in the system as well as the pairwise interaction potentials between them. In our case, the system was comprised of Mg^2+^ (or Ca^2+^), phosphate and water. The concentration of XP_2_ (where X = Mg^2+^ or Ca^2+^) is 1 mM. We used a 1-dimensional grid with a grid spacing of 0.025 Å and 131,072 grid points. Parameters for Mg^2+^ and Ca^2+^ were taken from the Amber force field that takes into account the charge-induced dipole interactions (17). We used the cSPC/E model for water, which introduces van der Waals terms for the hydrogen atoms of the SPC/E water model to prevent them to collapse in RISM (13). For phosphate, we used a single-site representation as in the coarse-grained model rather than an all-atom representation to overcome the convergence problems in RISM. To derive the phosphate parameters, we started with the Cl^−^parameters and tuned *ϵ* = 0.027 kcal/mol, and *R*_*min*_ = 2.6 Å (Eq. 6) to obtain the location of the first peak in the 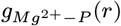 at around 2.5 Å (see Fig. 7B in the main text), which is somewhat longer than Mg^2+^–O distance 2.06 Å in the first hydration shell. Note that the position of the coarse-grained P site is located at the center of geometry of the phosphate group. We found that 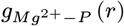 is not sensitive to the parameters provided the first peak is around 2.5 Å. We iteratively solved the RISM equations using the PSE-3 closure to a residual tolerance of 10^−12^ at 25°C. We emphasize that no simulation was performed at this stage. The pair distribution function between divalent ion–phosphate *g*_*X*_2+_*−P*_ (*r*) (X^2+^ is Mg^2+^ or Ca^2+^) is one of the direct outputs of the RISM program, from which we obtain 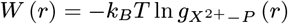.

#### 2. Development of coarse-grained TIS model

##### Coarse-grained RNA force field

Our goal is to produce an accurate coarse-grained model, which treats all the key interactions in a manner that can lead to quantitative predictions of thermodynamics and kinetics of RNA with arbitrary length. To this end, we build on the TIS model (18) in order to develop an RNA force field that treats divalent cations explicitly while describing the monovalent effects implicitly. Following our previous study (1, 19), each nucleotide is represented by three interaction sites, corresponding to phosphate (P), ribose (S) and base (B), where P and S represent the backbone; the B site depends on the nature of the nucleotide, and therefore carries the sequence information. The energy function has the following form:

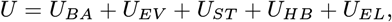

where *U*_*BA*_ is the *bonded term*, comprising of bond and angle restraints between connected beads. These constraints use harmonic potentials to keep the bonds and the angles close to the A-form helix. The parameters are the same as in our previous work (1, 19).

###### Excluded volume interactions *U*_*EV*_

We model *U*_*EV*_ using a modified LJ potential (1), which is evaluated using:

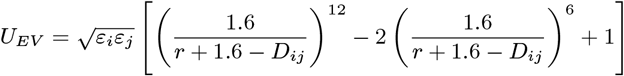

if *r ≤ D*_*ij*_ = *R*_*i*_ + *R*_*j*_. If *r > D*_*ij*_ then we set *U*_*EV*_ = 0. To allow for favorable base stacking, we set *D*_*BB*_ = 3.2 Å. It is not necessary to include excluded volume interactions between the divalent cation and P as it is taken into account in the effective potential displayed in Figs. 1 and 7A in the main text. The full set of parameters used in the simulations is given in Table S1.

**Table S1.**
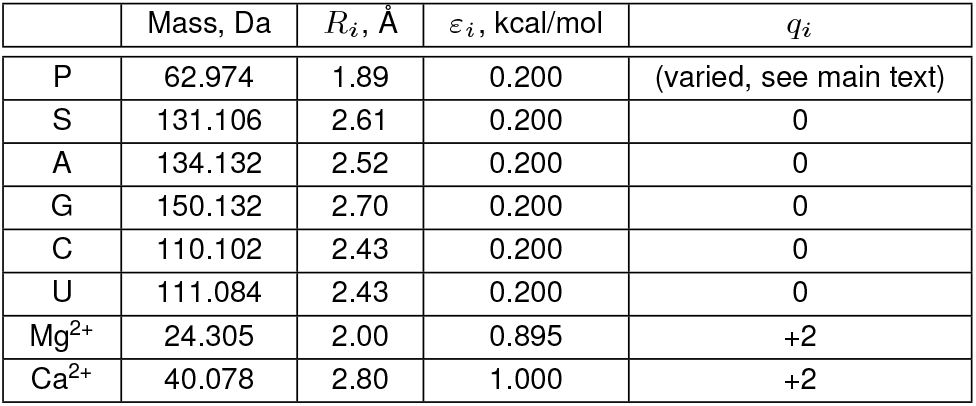
Parameters for excluded volume and Debye–Huckel interactions.

| | Mass, Da | $R_i$, Å | $\varepsilon_i$, kcal/mol | $q_i$ |
| --- | --- | --- | --- | --- |
| P | 62.974 | 1.89 | 0.200 | (varied, see main text) |
| S | 131.106 | 2.61 | 0.200 | 0 |
| A | 134.132 | 2.52 | 0.200 | 0 |
| G | 150.132 | 2.70 | 0.200 | 0 |
| C | 110.102 | 2.43 | 0.200 | 0 |
| U | 111.084 | 2.43 | 0.200 | 0 |
| Mg<sup>2+</sup> | 24.305 | 2.00 | 0.895 | +2 |
| Ca<sup>2+</sup> | 40.078 | 2.80 | 1.000 | +2 |

###### Justification for the divalent ion excluded volume parameters

The radius of the divalent cations used in this work to compute the excluded volume interactions are large (2.00 Å for Mg^2+^ and 2.80 Å for Ca^2+^) compared to the values used in atomistic simulations. In general, the size of various interaction sites has to be larger in coarse-grained models. We justify the value in Table S1 by arguing that the divalent ion radius in this work represents the fully hydrated form of the ion. For Mg^2+^, the relevant size is the radius of the hexahydrated form 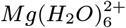, which coincides with the distance between Mg^2+^ and the oxygen atom in the first hydration shell 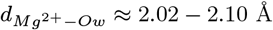. Therefore, in our model, we assume that Mg^2+^ ions do not dehydrate in order to interact with base or sugar moieties and other Mg^2+^ ions. Although the most frequent inner sphere coordination of Mg^2+^ occurs with the phosphate groups, it has been documented that Mg^2+^ also coordinates with nucleobases and less frequently with sugars (20). However, the neglect of such interactions, which likely do not contribute to charge neutralization of the phosphate groups, should not considerably affect the folding of RNA. From the perspective of folding thermodynamics, it is crucial to treat Mg^2+^–P interactions accurately, which we do rigorously using the RISM theory. We note that in our model, the divalent ion radius does not play any role in the divalent ion–phosphate interactions since these interactions are calculated based on the PMF (Eq. 5, main text). We do, therefore, allow the Mg^2+^ dehydration once they are near phosphate groups, as shown in Fig. 1 in the main text. The excellent predictions of the free energies for a variety of systems reported here show that it is crucial to account for the physics of divalent ion–phosphate interactions. This is accomplished using the RISM theory.

To ascertain that the radius of the divalent ion used here is physically reasonable, we compute the distance between the Mg^2+^ and sugars/bases in the simulations and compare them to the values in the crystal structure. The idea is to see if Mg^2+^ approaches the RNA in the simulations at distances that are too small or large. For the crystal structures, we took all RNA structures in the PDB that have Mg^2+^ and the resolution is at least 2.5 Å. With this criterion, we obtained 147 structures. We coarse-grained the RNAs and generated the histogram of the distances between Mg^2+^–base and Mg^2+^–sugar. Fig. S1 compares the radial distribution function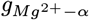 computed in the simulations of BWYV and the histogram generated from PDB. In the histogram, signals in the region *r <* 4.0 Å arise due to interactions with partially dehydrated Mg^2+^ ions, and therefore are not present in 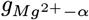. The key point here is that the closest Mg^2+^ could approach either moieties in the simulations is around 4.0 Å, which is in good agreement with the histogram, and therefore justifies the choice of the divalent ion radius used in our coarse-grained simulations. In addition, the agreement between our simulations and experimental data for a wide variety of thermodynamic properties furthermore justifies our choice of the divalent ion radius.

**Fig. S1.**
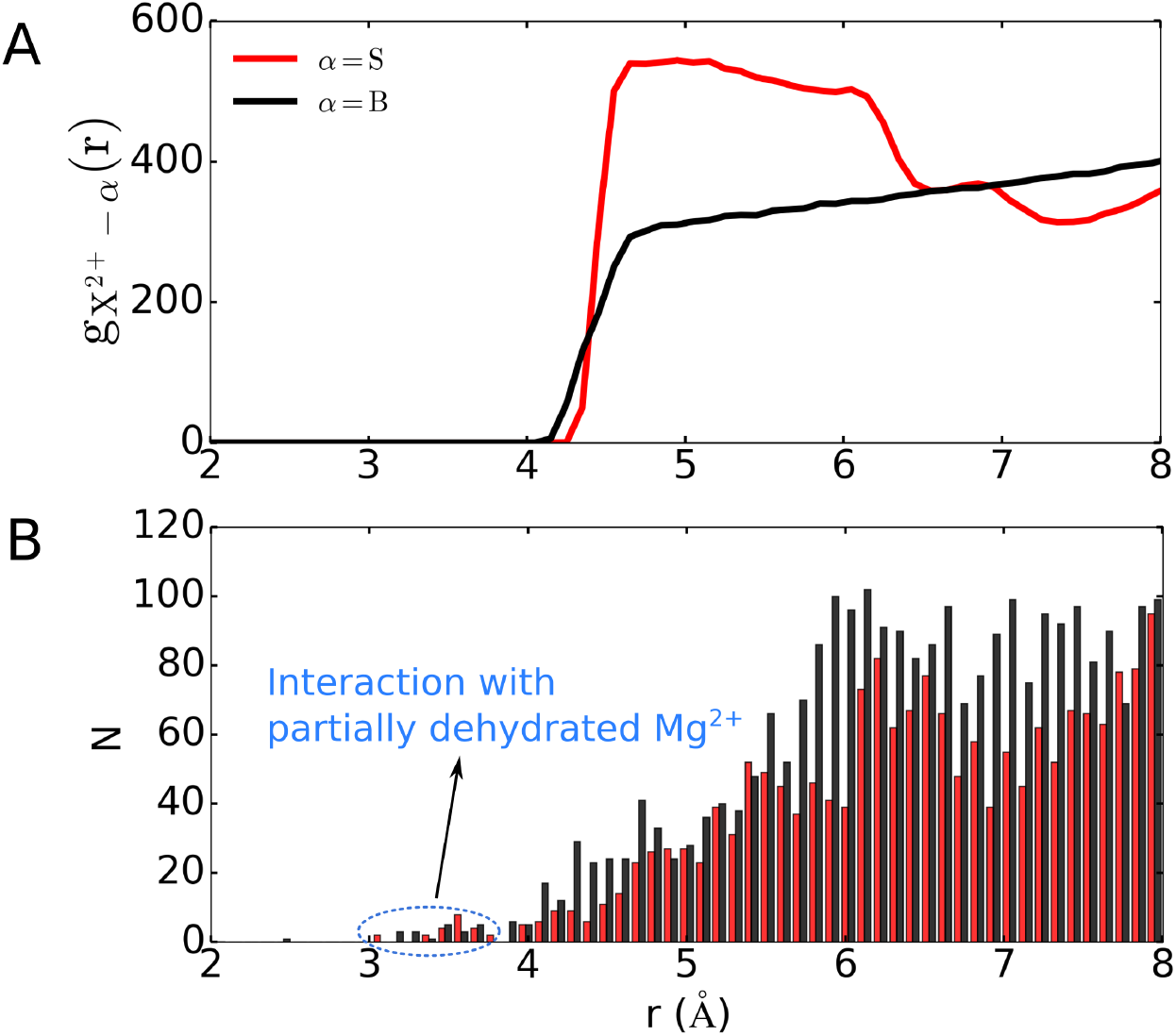
Comparison of the distances between Mg^2+^ and sugars (red) and bases (black). (A) Radial distribution function of Mg^2+^ and sugar (red) or base (black) calculated in simulations of BWYV. (B) Histogram of the distances between Mg^2+^ and sugar (red) or base (black) in 147 RNA structures. There is only a small occurrence of interaction between partially dehydrated Mg^2+^ with both sugar and base moieties in RNA at *r <* 4.0 Å.

###### Stacking interactions *U*_*ST*_

Interactions between two consecutive bases, or secondary stacking, are modeled using 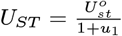 where *u*_1_ is a linear combination of harmonic constraints, which biases the stacking topology to the A-form helix (19), and 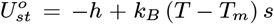 where *h* and *s* are independently obtained for the 16 nucleotide dimers by reproducing their experimental stacking thermodynamics (19). In the simulations, we computed the stability of the stacked dimers using:

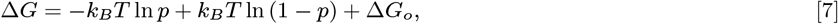

where *p* is the fraction of all sampled conformations for which *U*_*ST*_ < *−k*_*B*_*T*. The parameters *h* (but not *s*) in 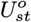, thus, are functions of a single free energy correction term, t:, *ΔG*_*o*_. The value of t:, *ΔG*_*o*_, assumed to be a constant for all dimers, is used to adjust the balance between stacking and hydrogen bonding, which is essential to accurately reproduce RNA thermodynamics (see below).

Stacking between non-consecutive bases is detected from the input structure using the geometric criteria reported elsewhere (21). We evaluated tertiary stacking using 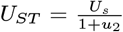, where *U*_*s*_ = *−*5.0 kcal/mol, and *u*_2_ is also a linear combination of harmonic constraints, similarly to *u*_1_, but instead is chosen to bias the stacking topology to the crystal structure. Since a base could stack with others on both sides, we keep track of both the number of stacking each side participates in during the simulations. Each side is allowed to stack with a maximum of two other bases, and tertiary stacking is given a higher priority over secondary stacking. In other words, once tertiary stacks are formed (the two bases are closer than 10.0 Å) and the side reaches the maximum stacking capacity then the secondary stacking is disallowed.

###### Hydrogen bond potential *U*_*HB*_

We used 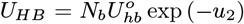 where *u*_2_ has the same form as tertiary stacking, biasing the structure towards an A-form RNA for canonical bonds (G-C, A-U and G-U) or the experimental structure for non-canonical bonds; *N*_*b*_ is the number of hydrogen bonds between the beads. For Watson–Crick base pairing, *N*_*b*_ is 2 for A-U and G-U, and 3 for G-C. For non-canonical bonds, *N*_*b*_ is computed from the experimental structure of the RNA. Hydrogen bonds involving S and P beads are also considered. We use 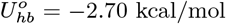, which is fit in order to reproduce heat capacities of RNA hairpins and pseudoknots (Fig. S2). Our model also permits non-native base pairing formed between G and C, A and U, G and U separated by at least 4 nucleotides along the chain. Thus, the folded structures can be disrupted, allowing for non-native structures to be populated in the simulations. Each bead has a maximum number of hydrogen bonds it could potentially form, and one base is involved in only one canonical base-pair.

###### Choice of *ΔG*_*o*_ and 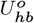

Following our previous studies, we calibrated the parameters to reproduce known experimental quantities. Here, we adjusted t:, *ΔG*_*o*_ in Eq. 7 (0.90 kcal/mol) and 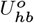 (−2.70 kcal/mol) to fit the heat capacity of human telomerase RNA hairpin (hTR HP) in 200 mM KCl and the Beet Western Yellow Virus pseudoknot (BWYV PK) in 500 mM KCl. In Fig. S2, the values of the melting temperatures for both the hTR HP and BWYV PK predicted theoretically are in good agreement with experiments.

**Fig. S2.**
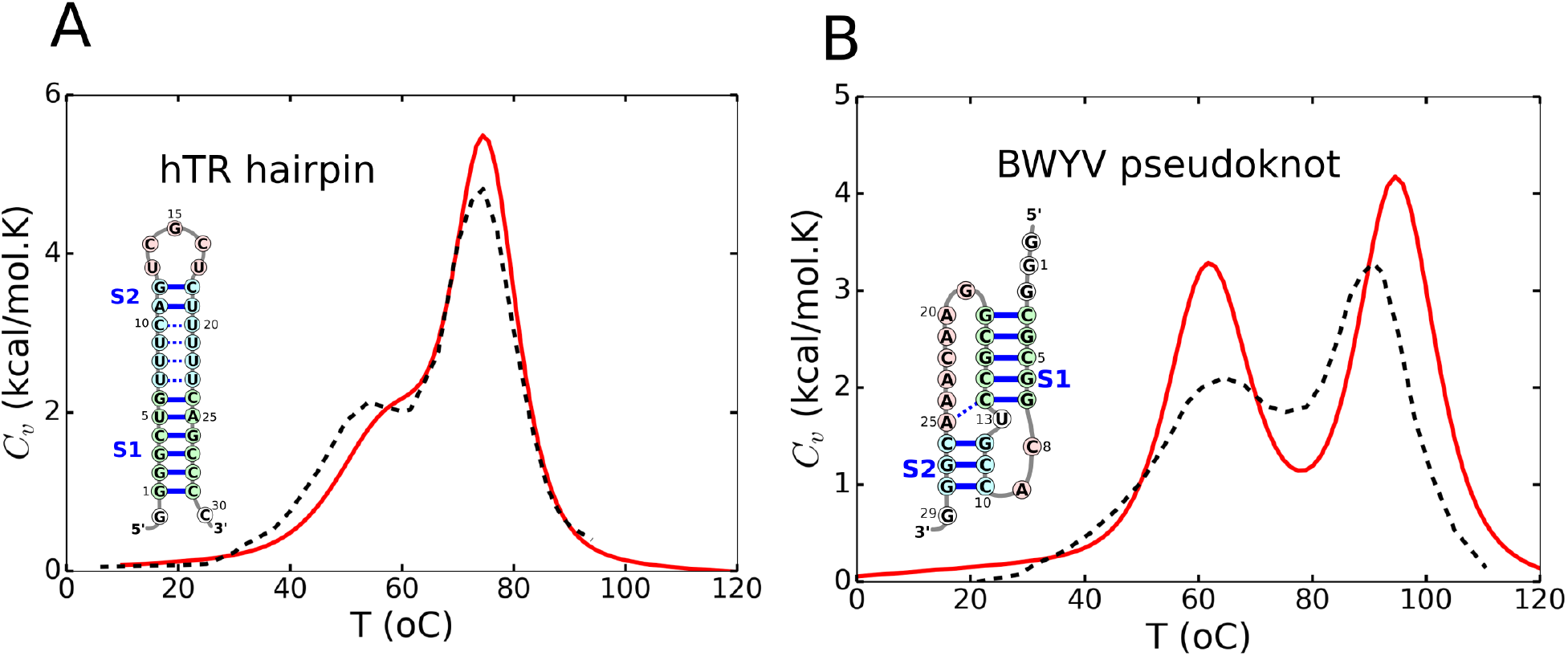
Heat capacity of hTR hairpin (A) and BWYV pseudoknot (B). Simulations were performed for hTR in 200 mM KCl and for BWYV in 500 mM KCl at pH=7.0 with no divalent cations. Experimental data for hTR and BWYV are black dashed lines, taken from Ref. (22) and (23), respectively. Simulation data are shown as red lines.

###### Electrostatic interactions *U*_*EL*_

We developed a new way to treat the interaction of ions with a charged phosphate group. A typical buffer used in RNA folding experiments (Tris for example) contains known amount of monovalent ions. To this buffer, a solution containing divalent cations is added in order to induce RNA folding. Thus, in such experiments, the solution contains a mixture of divalent and monovalent ions. Treating both the ions on equal footing, which we previously carried out for a few RNA constructs including the *Azoarcus* ribozyme (1), is computationally demanding. As explained here and in the main text, we treated monovalent ions implicitly and Mg^2+^/Ca^2+^ explicitly. This procedure is justified because divalent cations are indispensable for the folding of most RNAs, with the exception of some small hairpins and pseudoknots. It is, therefore, important to include divalent ions explicitly to take into account the ion size and their specific interactions with the RNA. On the other hand, the majority of monovalent ions interacts with the RNAs via non-specific electrostatic interactions, screening the charge–charge repulsion between phosphate groups. There are few examples where specific interactions between monovalent ions and RNA play an essential role in RNA folding (24–26). Hence, it is reasonable to treat the monovalent ion as a continuum using classical Debye–Huckel theory. In most RNA folding experiments, the concentration of monovalent ions in the buffer solution typically is in far excess of the divalent cation, and therefore they screen the interactions between the divalent ions. Thus, we assume that the Debye–Huckel potential accurately describes the electrostatic interactions between divalent cations and P–P repulsions. With this approximation, *U*_*EL*_ is the sum of pairwise interactions between all the divalent cations, repulsions between the P groups, and attractive interactions between the divalent cations and the P groups. The bare charge on P is replaced by an effective charge, *Q* (*T, C*_1_, *C*_2_), which is calculated using the counter ion condensation theory. The value of *Q* (*T, C*_1_, *C*_2_) depends on the temperature, *T*, as well as the concentrations of monovalent (*C*_1_) and divalent cations (*C*_2_). It only remains to determine the interaction between the divalent cations and P, which is given by Eq. 5 in the main text. The PMF, *W* (*r*), is calculated using the RISM theory in liquid state physics, described in the previous section.

##### Validation of the ion condensation theory

The assumption of the Oosawa–Manning counter ion condensation theory is that ions at distances that are larger than the size of the RNA corresponding to the bulk (B), and the ones condensed (C) onto the polyanion are at equilibrium (27). The value of the renormalized charge on the phosphate group is calculated by equating the chemical potentials of the B and C ions. In the main text, we derived an approximate expression of charge neutralization to obtain the effective charge on the phosphates using this physical picture. In order to assess if our estimate of charge renormalization is reasonable, we rely on experimental techniques that probe both the monovalent and divalent ion atmosphere around nucleic acids such as ion counting (BE-AES) or anomalous small-angle X-ray scattering (ASAXS) (28–30). Fig. S3 compares *θ*_*Na*_^+^ and 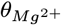, 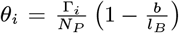, computed using the theory at 20 mM NaCl and results from ion counting experiment for a 24bp duplex DNA. The experimental *θ* are calculated using the assumption that the total DNA charge neutralized is similar to the theory, hence the factor 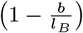. This is necessary because we set *b* = 4.4 Å in our theory and simulations. We had to choose DNA because the data for RNA is currently not available. Because of the physics of ion condensation likely does not depend greatly on the differences between DNA and RNA, comparison with duplex DNA is sufficient to validate our theory. The calculated values are in good agreement with experiments, indicating that our theory describes ion competition in a buffer containing both monovalent and divalent cations accurately.

**Fig. S3.**
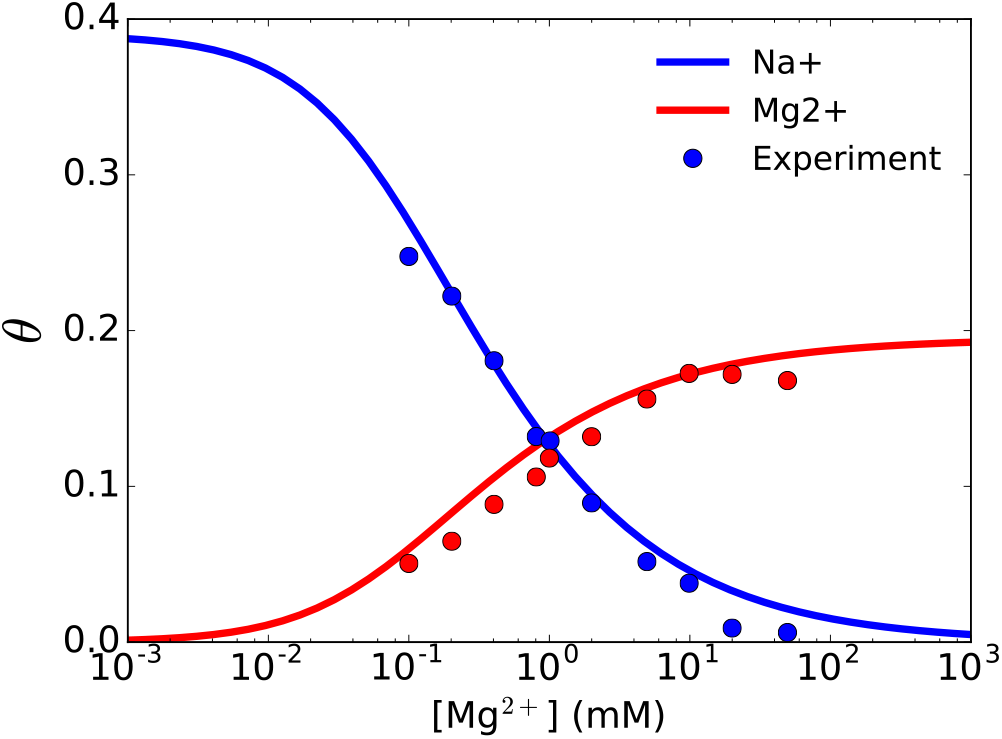
Competition between Na^+^ and Mg^2+^ ions around a 24bp DNA duplex. The plot shows the number of condensed ions per phosphate group at a fixed 20 mM NaCl solution as a function of Mg^2+^ concentration. Calculations were done by analytically solving Eqs. 3 and 4 (main text). The blue and red circles are experimental data from Ref. (28) and 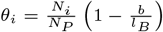, with *b* = 4.4 Å and *N*_*P*_ = 46 is the total number of phosphate groups in the DNA.

#### 3. Simulation details and data analyses

##### Simulations

We performed simulations using the Langevin dynamics with the CG force field using an in-house code, which is available at https://github.com/tienhungf91/RNA_cg. Divalent cations were randomly added to a cubic box containing an RNA molecule, whose initial coordinates were taken from the structure of the folded state in the PDB. The box size varied from 700-3,000 Å depending on the bulk concentration of divalent cations. We used large boxes to make sure that at least 200 divalent cations were present in the simulations. Enlarging the box size does not introduce more particles into the system, but only dilutes the divalent cation concentration. The performance of our model, therefore, is insensitive to the box size, allowing us to probe the effect of the arbitrarily small concentration of divalent cations on RNA folding. This is another advantage of treating monovalent ions implicitly. We used periodic boundary conditions in the simulations to minimize the effect of finite box size. Numerical integration of the equations of motion was performed using the leap-frog algorithm with the time step *h* = 0.05*⋅* where 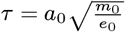 is the unit of time, *a*_0_ = 1 Å, *m*_0_ = 1 Da and *e*_0_ = 1 kcal/mol. We performed low-friction dynamics to increase the sampling efficiency of the conformations, in which the viscosity of water was reduced 100 times (31). Snapshots were recorded every 10,000 steps, from which only the last two-thirds were used to compute all the quantities of interest.

##### Calculation of the heat capacity

We performed replica-exchange simulation (REMD) at several temperatures (32). Exchange was attempted every 5,000 steps between neighboring replicas. The system energy was recorded every 10,000 steps, and the heat capacity was computed using 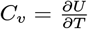 with WHAM. The REMD was found to give converged results after ~ 5 × *θ* 10^8^ integration steps.

##### Calculation of the folding free energy, *ΔG* (*c*)

For a given monovalent concentration *C*_1_, the folding free energy of the RNA is calculated using: (19)

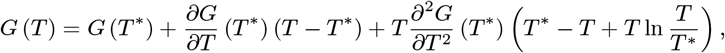

where *T\** is the reference temperature. At temperatures that are low compared to the melting temperature, only the folded state of the RNA is predominantly populated. Thus, the free energy of the folded state, *G*_*f*_ (*T*), can be determined by using, for instance, *T \** = 10°C as the reference temperature. Similarly, the free energy of the unfolded state, *G*_*u*_ (*T*), is computed using *T \** = 120°C. The free energy of the intermediate state, *G*_*i*_ (*T*), is computed with *T\** in between the two melting temperatures. For BWYV PK, *T \** = 70°C (see Fig. S2). We performed REMD simulations at several temperatures and used WHAM to compute *G* (*T*). The free energy for each state is then calculated using the above equation with appropriate reference temperatures. The folding free energy is then evaluated using *ΔG*_*f−u*_ (*T*) = *G*_*f*_ (*T*) *− G*_*u*_ (*T*).

##### Equilibrium between the bulk and condensed ions and Γ_*X*_2+

Because the divalent ions are attracted strongly to the highly negatively charged RNA, the actual ion bulk concentration differs from the concentration computed by dividing the total number of ions by the volume of the simulation box. Failure to account for the equilibrium between these populations results in an incorrect calculation of Γ_*X*_2+ and *ΔG*_*X*_2+ _*−RNA*_. One could enlarge the simulation box to alleviate this problem, but it is computational demanding. Instead, we calculated the ion concentration in the bulk *C*_2_ after the concentration profile of ions, *C*_2_ (*r*), plateaus at large separation from the RNA (Fig. S4). The preferential interaction coefficient is then evaluated as 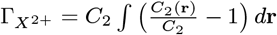. In practice, we truncated the integration at the distance where *C*_2_ (*r*) = *C*_2_.

**Fig. S4.**
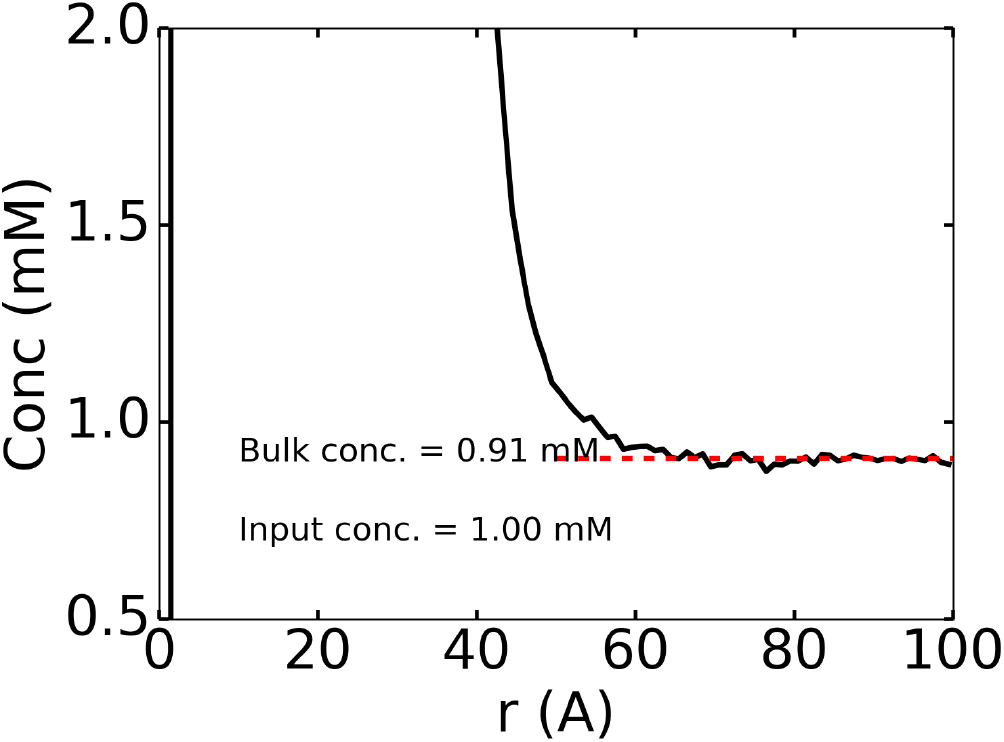
Concentration profile of Mg^2+^ around adenine riboswitch. The bulk concentration is computed at a large separation from the center-of-mass of the RNA and subsequently used for *Γ*_*Mg*_2+ determination.

##### X^2+^–RNA free energy

The plot of Γ_*X*_2+ vs. ln *C*_2_ curve (in Fig. 2 in the main text, for example) (X^2+^ is either Mg^2+^ or Ca^2+^) is fit using a fourth order polynomial, *y* = *b* (*x − a*)^2^ + *c*(*x − a*)^3^ + *d*(*x − a*)^4^ and the fit polynomial is integrated analytically, which allows us to evaluate the integral in Eq. 7 analytically and obtain *ΔG*_*X*_2+, _*RN A*_.

##### Fraction of native contacts

The fraction of native contacts, *Q* (*X*_*k*_), for an RNA conformation *X*_*k*_ is computed using: (33)

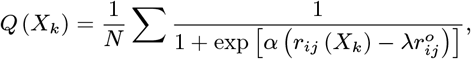

where the sum runs over *N* pairs of native contacts (i,j) separated by 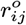 in the crystal structure, *α* = 5 Å-1 is a smoothing parameter and *λ* = 1.5 accounts for fluctuations when contacts formed. The list of *N* contacts are determined by native hydrogen bonds and stackings in the PDB structure, including secondary and tertiary interactions.

#### 4. Robustness of the model

##### BWYV

In order to provide additional evidence of the robustness of the force field, we calculated the free energy difference, *ΔG*_*F −U*_ (*c*), between the folded and unfolded states, as a function of monovalent salt concentrations *C*_1_ for BWYV pseudoknot. Fig. S5 shows that the simulated values of *ΔG* (*c*) are in very good agreement with experiments.

**Fig. S5.**
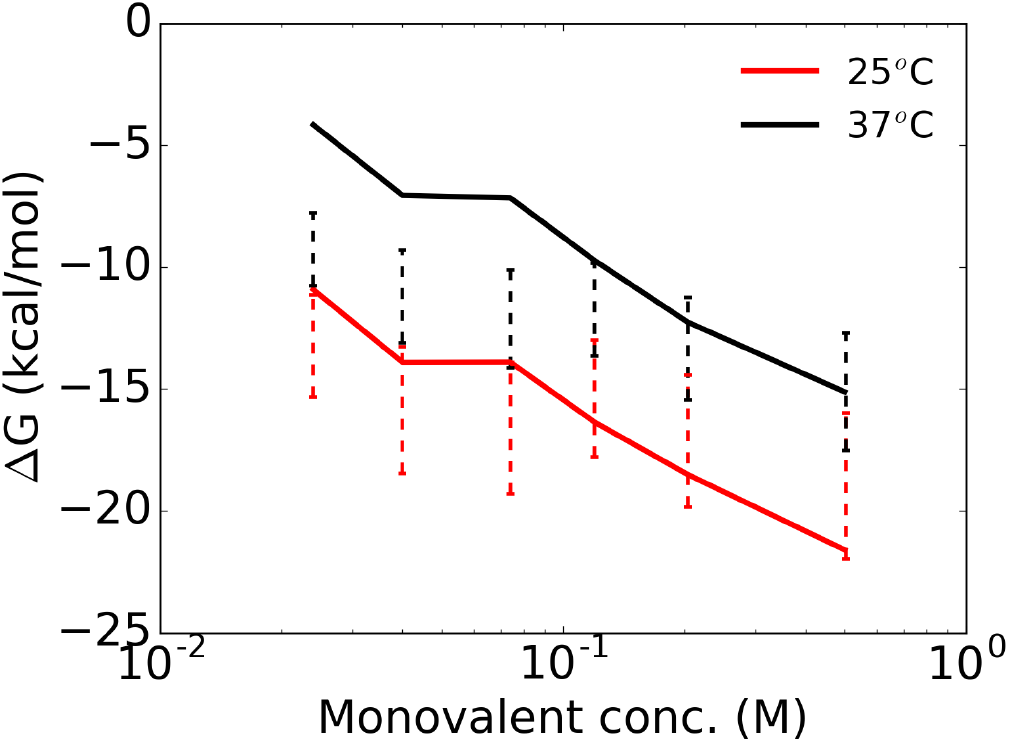
Free energy difference (*ΔG*) between the folded and unfolded states of BWYV pseudoknot as a function of monovalent concentration (*C*_1_) evaluated at 25°C (red) and 37°C (black). Calculated values are plotted as solid lines. The folding enthalpy and entropy reported in Ref. (34) were used to compute the experimental free energies shown with error bars. The calculated values of *ΔG* from simulations are in remarkable agreement with experiments, except for an underestimation at 37°C at low ion concentrations.

##### 58-nt rRNA

As a further validation, we calculated the heat capacity (*C*_*v*_) of the 58-nt fragment of rRNA (Fig. S6) and compared the results with the UV absorbance data (35). The melting temperature at 20 mM KCl, identified with the maximum in *C*_*v*_, agrees reasonably with the experimental data. The value of the high temperature peak in *C*_*v*_ at 60 mM KCl and 1 mM MgCl_2_ is in good agreement with the experimental data. However, the shoulder at the lower temperature obtained in the simulations, if it exists at all, is much less pronounced in experiments. The results in Figs. S2, S5 and S6 show that our RNA force field is sufficiently accurate to reproduce many aspects of the thermodynamics for several RNAs over a wide range of ion concentrations. It is worth remarking that currently there is no other computational model that can calculate ion-dependent folding thermodynamic properties of RNA, such as free energy changes and heat capacities, let alone achieve the level of accuracy reported here.

**Fig. S6.**
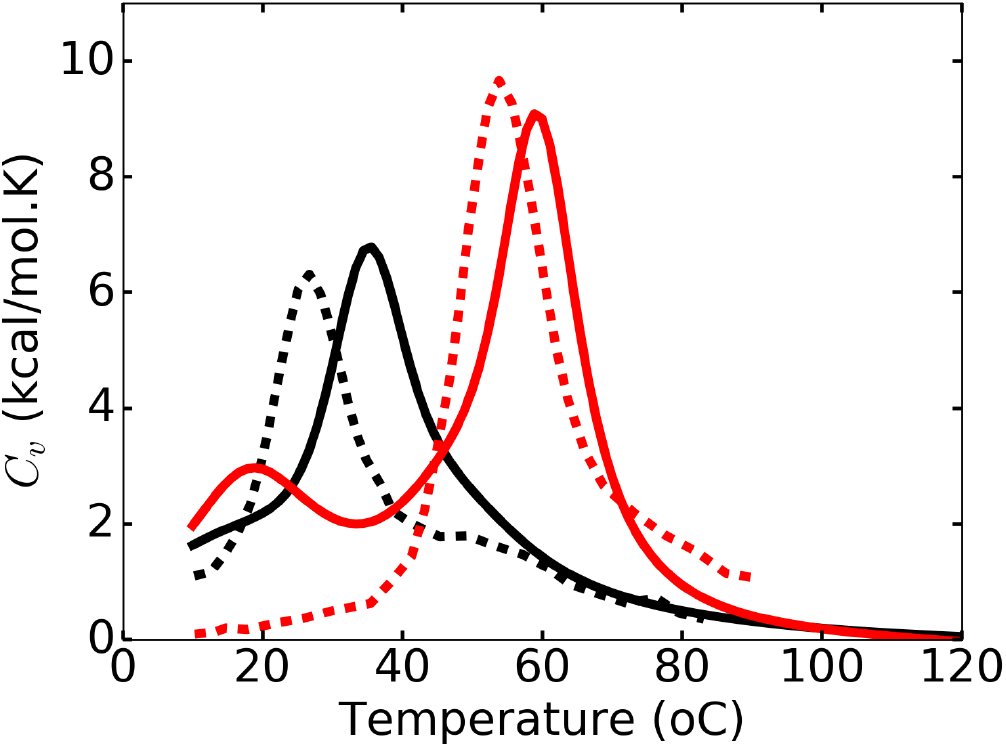
Heat capacity of the 58-nt rRNA at two salt concentrations. Black–20 mM KCl (no Mg^2+^), red –60 mM KCl + 1 mM MgCl_2_. Dashed curves show UV absorbance data from experiments (35), solid lines are from simulations. We should stress that UV absorbance data is not the same physical variable as *C*_*v*_, which is computed using the fluctuations in energy. Therefore, for the purposes of comparison, only the peak positions are relevant. The differences in the major melting temperature between the simulations and experiments are ~ 9°C at 20 mM KCl (no Mg^2+^) and ~ 5°C at 60 mM KCl + 1 mM MgCl_2_. The comparison further demonstrates that the agreement between simulations and experiments is excellent.

**Fig. S7.**
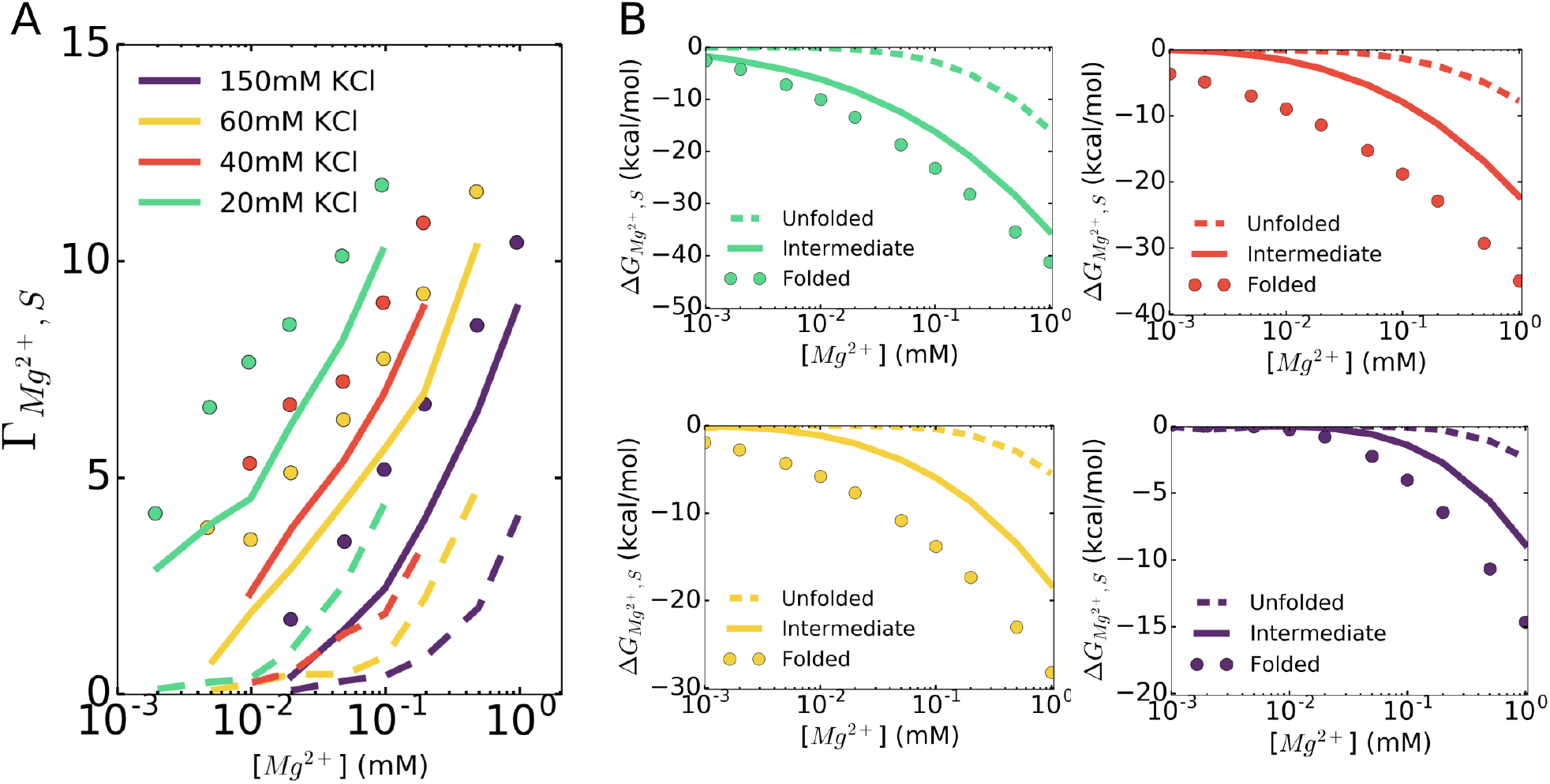
(A) Γ_*Mg*_2+, _*S*_ and (B) *ΔG*_*Mg*_2+,_*S*_ for the folded, intermediate and unfolded states of 58-nt rRNA. Simulations were performed by constraining the ensemble of RNA conformations to specific states. See the main text for details.

**Fig. S8.**
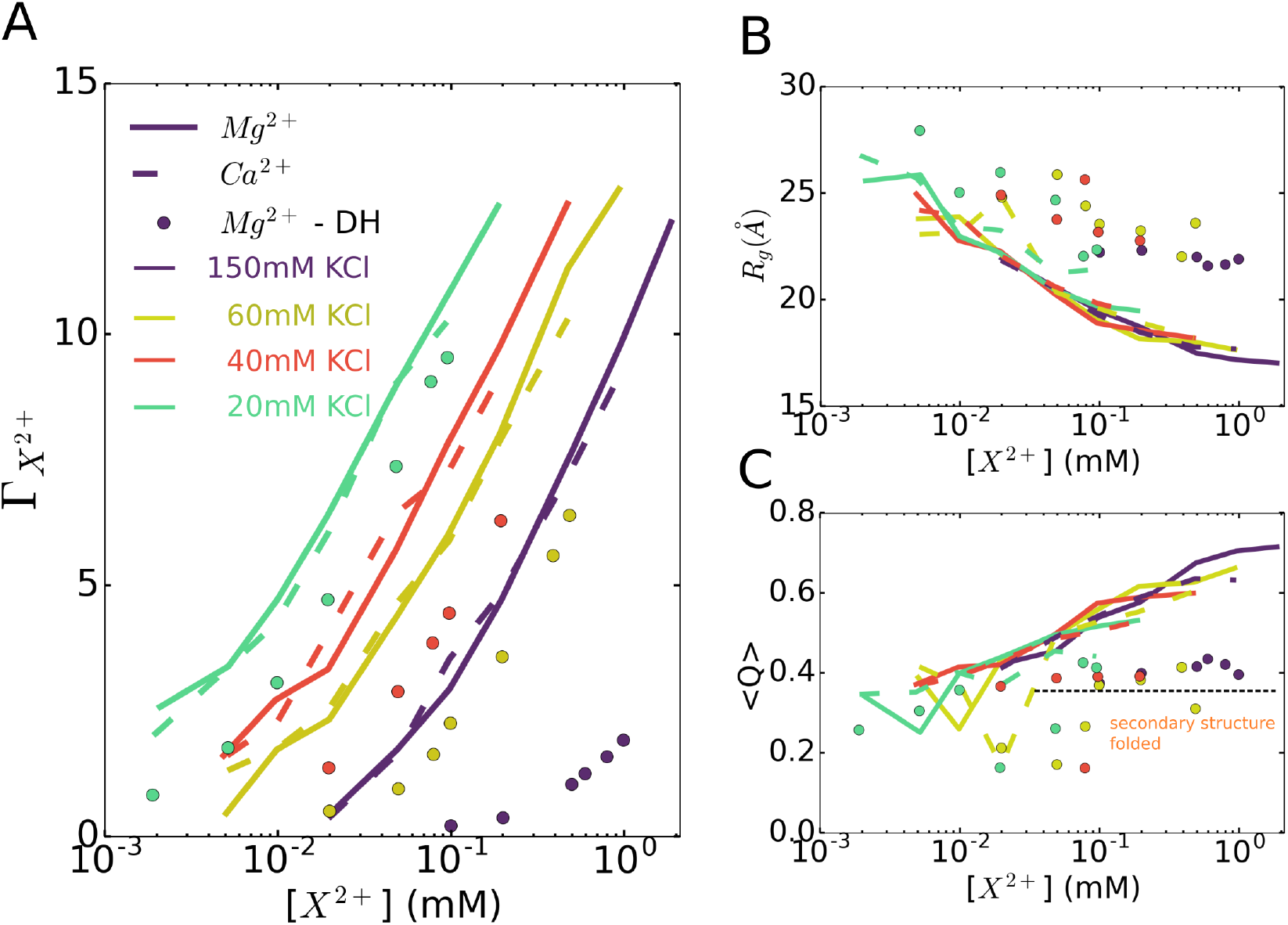
Shown here are the full data for Fig. 7 in the main text for the 58-nt rRNA.

**Fig. S9.**
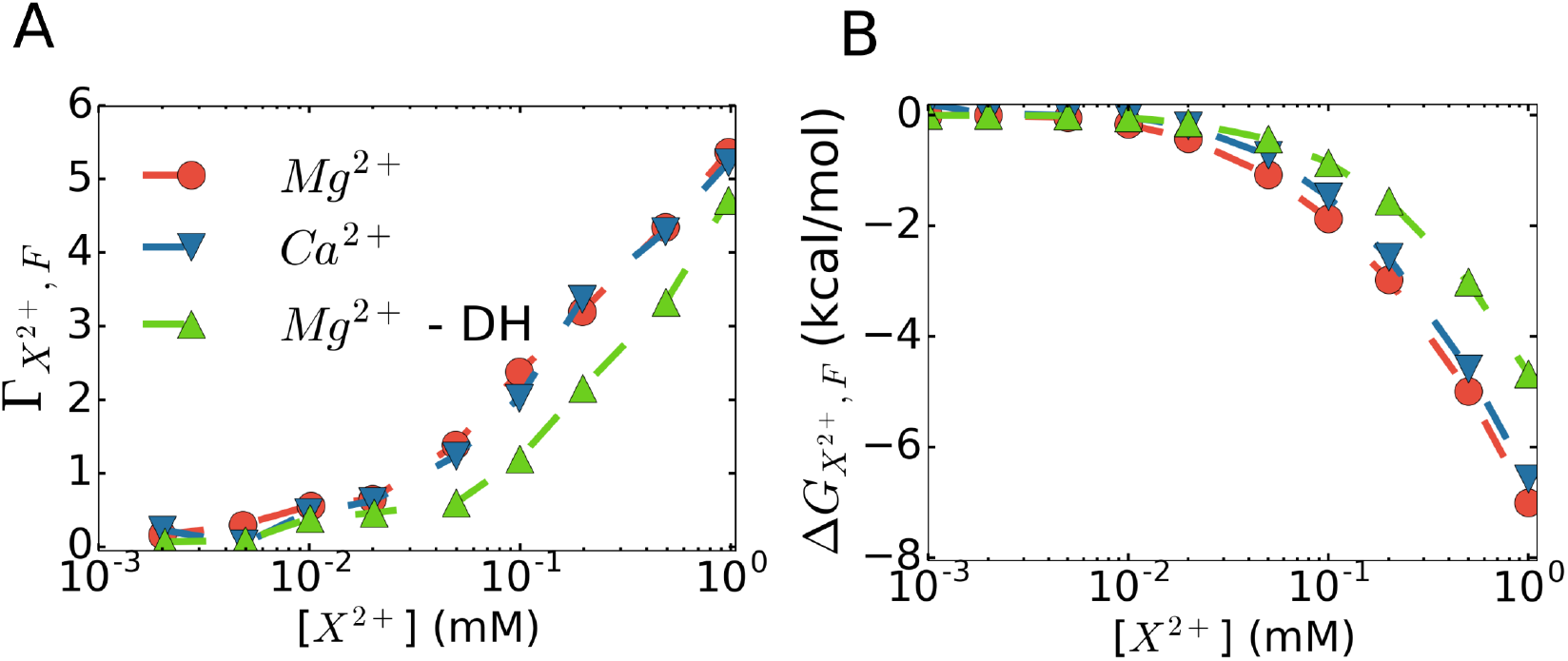
Ion preferential interaction coefficients (A) and free energies of divalent ion–RNA interactions (B) of Mg^2+^ (red) and Ca^2+^ (blue) computed for BWYV at 54 mM KCl. Calculations were also performed for Mg^2+^, in which the interactions are treated at the Debye–Huckel level (green).

**Fig. S10.**
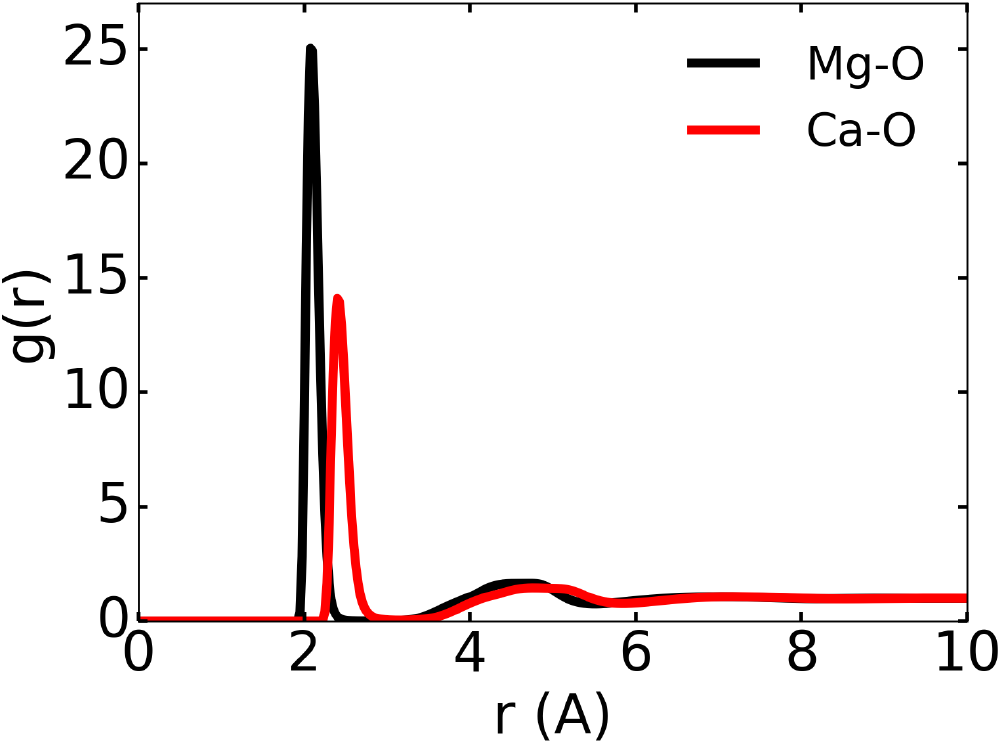
g(r) between divalent ion and water computed from RISM theory.

